# The ribosomal RNA m^5^C methyltransferase NSUN-1 modulates healthspan and oogenesis in *Caenorhabditis elegans*

**DOI:** 10.1101/2020.03.16.993469

**Authors:** Clemens Heissenberger, Teresa L. Krammer, Jarod A. Rollins, Fabian Nagelreiter, Isabella Stocker, Ludivine Wacheul, Anton Shpylovyi, Santina Snow, Johannes Grillari, Aric N. Rogers, Denis L.J. Lafontaine, Markus Schosserer

**Affiliations:** Department of Biotechnology, Institute of Molecular Biotechnology, University of Natural Resources and Life Sciences, Vienna, Muthgasse 18, 1190 Vienna, Austria; MDI Biological Laboratory, 159 Old Bar Harbor Rd, 04609 Maine, USA; RNA Molecular Biology, Fonds de la Recherche Scientifique (F.R.S./FNRS), Université Libre de Bruxelles (ULB), B-6041 Charleroi-Gosselies, Belgium; Christian Doppler Laboratory for Biotechnology of Skin Aging, Department of Biotechnology, BOKU, Vienna, Austria; Ludwig Boltzmann Institute of Experimental and Clinical Traumatology, 1200 Vienna, Austria

## Abstract

Our knowledge about the repertoire of ribosomal RNA modifications and the enzymes responsible for installing them is constantly expanding. Previously, we reported that NSUN-5 is responsible for depositing m^5^C at position C2381 on the 26S rRNA in *Caenorhabditis elegans*.

Here, we show that NSUN-1 is writing the second known 26S rRNA m^5^C at position C2982. Depletion of *nsun-1* or *nsun-5* improved locomotion at midlife and resistance against heat stress, however, only soma-specific knockdown of *nsun-1* extended lifespan. Moreover, soma-specific knockdown of *nsun-1* reduced body size and impaired fecundity, suggesting non-cell-autonomous effects. While ribosome biogenesis and global protein synthesis were unaffected by *nsun-1* depletion, translation of specific mRNAs was remodelled leading to reduced production of collagens, loss of structural integrity of the cuticle and impaired barrier function.

We conclude that loss of a single enzyme required for rRNA methylation has profound and highly specific effects on organismal physiology.

## Introduction

Ageing is a complex biological process, characterized by progressive aggravation of cellular homeostasis defects and accumulation of biomolecular damages. According to the ‘disposable soma theory’ of ageing, organisms may invest energy either in reproduction or in somatic maintenance (Kirkwood and Holliday, 1979). This explains why most lifespan-extending interventions come at the cost of decreased fecundity. D*e novo* protein synthesis by ribosomes, the most energy-demanding process in living cells, affects the balance between ageing and reproduction. In fact, reduced overall protein synthesis was shown to extend lifespan in several model organisms, including the nematode *Caenorhabditis elegans* (Hansen et al., 2007; Pan et al., 2007; Syntichaki et al., 2007). Although some evidence indicates that a link between ribosome biogenesis and gonadogenesis in *C. elegans* may exist (Voutev et al., 2006), the precise relationship between these pathways in multicellular organisms is still poorly understood. It is conceivable that the optimal function of ribosomes, which requires the presence of ribosomal RNA (rRNA) and ribosomal protein (r-protein) modifications, is monitored by the cell at several stages during development. Thus, introduction of these modifications might participate in the control of cell fate and cell-cell interactions during development (Hokii et al., 2010; Voutev et al., 2006).

Eukaryotic ribosomes are composed of about 80 core r-proteins and four different rRNAs, which together are assembled into a highly sophisticated nanomachine carrying the essential functions of mRNA decoding, peptidyl transfer and peptidyl hydrolysis (Ban et al., 2014; Natchiar et al., 2017; Penzo et al., 2016; Sharma and Lafontaine, 2015; Sloan et al., 2017). Until recently, ribosomes were considered as static homogenous ribonucleoprotein complexes executing the translation of cellular information from mRNA to catalytically active or structural proteins. However, mounting evidence suggests the possibility of ribosomes being heterogeneous in composition with the possibility that some display differential translation with distinct affinity for particular mRNAs (Genuth and Barna, 2018). Such heterogeneity in composition may originate from the use of r-protein paralogs, r-protein post-translational modifications, or rRNA post-transcriptional modifications. Indeed, around 2% of all nucleotides of the four rRNAs are decorated with post-transcriptional modifications, which are introduced by specific enzymes such as dyskerin and fibrillarin, guided by specific small nucleolar RNAs (snoRNA) (Penzo et al., 2016; Sloan et al., 2017). The most abundant rRNA modifications are snoRNA-guided 2′O-methylations of nucleotide ribose moieties and isomerization of uridine to pseudouridine (Ψ). However, some base modifications, which occur less frequently than 2’O-methylations of ribose and pseudouridines, are installed by specific enzymes, which were largely assumed to be stand-alone rRNA methyltransferases. One exception is the acetyltransferase Kre33 (yeast)/NAT10 (human), which is guided by specialized box C/D snoRNPs (Sharma et al., 2015; Sleiman and Dragon, 2019). Most of these base modifications are introduced at sites close to the decoding site, the peptidyl transferase centre, or the subunit interface. Intriguingly, prokaryotes and eukaryotes share the majority of modifications located in the inner core of the ribosome (Natchiar et al., 2017).

In eukaryotes inspected so far, the large ribosomal subunit contains two m^5^C residues. This is notably the case in budding yeast (on 25S rRNA), in the nematode worm (26S) and in human cells (28S) (Sharma and Lafontaine, 2015). Rcm1/NSUN-5, an enzyme of the NOP2/Sun RNA methyltransferase family, is responsible for introducing m^5^C at residue C2278 and C2381 on 25S/26S rRNA in yeast and worms, respectively (Gigova et al., 2014; Schosserer et al., 2015; Sharma et al., 2013). Recently, our group and others identified the conserved target cytosines in humans and mice, namely C3782 and C3438, respectively (Janin et al. 2019, Heissenberger et al. 2019). We also reported that lack of this methylation is sufficient to alter ribosomal structure and ribosome fidelity during translation, while extending the lifespan and stress resistance of worms, flies and yeast (Schosserer et al., 2015). However, the identity of the second worm m^5^C rRNA methyltransferase remains unknown.

Here, we report that NSUN-1 is responsible for writing the second *C. elegans* 26S m^5^C (position C2982). We then investigate the physiological roles of NSUN-1, comparing them systematically to those of NSUN-5. We show that NSUN-1 and NSUN-5 distinctly modulate fundamental biological processes such as ageing, mobility, stress resistance and fecundity. In particular, depletion of *nsun-1* impairs fecundity, gonad maturation and remodels translation of specific mRNAs leading to cuticle defects. We conclude that loss of NSUN-1 introducing a single rRNA modification is sufficient to profoundly and specifically alter ribosomal function and, consequently, essential cellular processes.

## Results

### NSUN-1 is responsible for writing m^5^C at position C2982 on *C. elegans* 26S rRNA

Previously, we showed that an m^5^C modification is introduced at position C2381 on the 26S rRNA of *C. elegans* large ribosomal subunit by NSUN-5 (Adamla et al. 2019; Schosserer et al. 2015), which is required to modulate animal lifespan and stress resistance (Schosserer et al., 2015). On this basis, we were interested to learn if other related rRNA methyltransferases in *C. elegans* might display similar properties.

Therefore, we investigated the RNA substrate of NSUN-1 (also formerly known as NOL-1, NOL-2 or W07E6.1) and its potential roles in worm physiology. NSUN-1 is also a member of the NOP2/Sun RNA methyltransferase family. Since there are only two known m^5^C residues on worm 26S rRNA (Sharma and Lafontaine, 2015; Trixl and Lusser, 2019), one of them at C2381, being installed by NSUN-5, we speculated that NSUN-1 might be required for introducing the second m^5^C residue at position C2982. Notably, both 26S m^5^C sites are localized close to important functional regions of the ribosome, and they are highly conserved during evolution (Fig. 1A,B).

**Figure 1:**
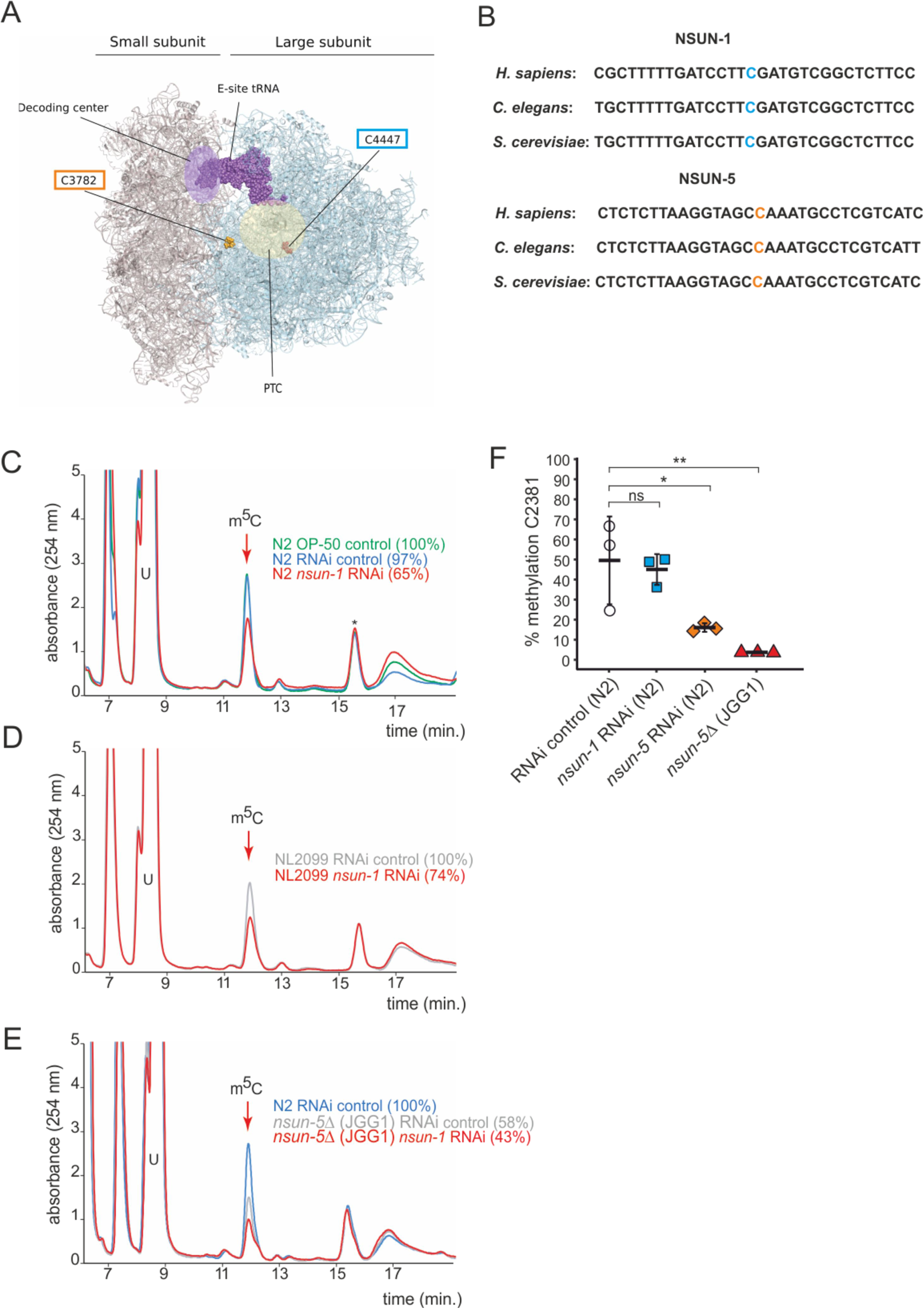
NSUN-1 is responsible for large ribosomal subunit 26S rRNA m^5^C methylation. **A,** Location of the two eukaryotic large ribosomal subunit m^5^C residues within the 3D structure of the human ribosome. For reference, important functional sites are indicated (DCS = decoding site, PTC = peptidyl transferase centre). In *C. elegans*, NSUN-1 is responsible for m^5^C2982 (this work) while NSUN-5 installs m^5^C2381 (Schosserer et al. 2015). **B,** Regions surrounding the sites modified by NSUN-1 and NSUN-5 are evolutionarily conserved between yeast, worms and humans. The modified cytosine is indicated. **C-E,** Purified 26S rRNA was isolated by sucrose gradient centrifugation, digested to single nucleotides and analysed by quantitative HPLC. *nsun-1* knockdown consistently leads to a decrease of m^5^C levels. **C,** N2 worms were analysed as either: untreated (OP-50), treated with an RNAi control or with a *nsun-1* targeting RNAi. **D,** NL2099 RNAi hypersensitive worms were treated with the RNAi control or with the *nsun-1* targeting RNAi. **E,** N2 strain treated with RNAi control and the *nsun-5* deletion strain (JGG1) treated with control RNAi or a *nsun-1* targeting RNAi. For quantification of m^5^C peak area, the peak was normalized to the peak eluting at 16 min (asterisk). **F,** Quantification of the enzymatic activity of NSUN-5 using the COBRA assay for N2 worms, subjected to either *nsun-5* or *nsun-1* RNAi, and the *nsun-5* mutant strain JGG1 (*nsun-5*Δ). Loss of *nsun-5* leads to significantly decreased methylation levels at C2381, whereas *nsun-1* RNAi does not alter methylation at this site (three independent biological replicates, one-way ANOVA with Dunnett′s post test, α=0.05, *P<0.05, **P<0.01).

In order to test if NSUN-1 is involved in large ribosomal subunit m^5^C methylation, 26S rRNA was purified from worms treated with siRNAs specific to NSUN-1-encoding mRNAs on sucrose gradients, digested to single nucleosides and analysed by quantitative HPLC. In our HPLC assay, the m^5^C nucleoside eluted at 12 min, as established with a m^5^C calibration control (data not shown).

The depletion of NSUN-1 was conducted in two genetic backgrounds: N2 (wildtype), and NL2099 (an RNAi-hypersensitive strain due to mutation in *rrf-3*) (Figure 1–figure supplement 1). Treating N2 worms with an empty control vector, not expressing any RNAi, did not significantly reduce the levels of 26S rRNA m^5^C methylation (Fig. 1C, 97% instead of 100%). Interestingly, treating N2 worms with an RNAi construct targeting *nsun-1* led to a reduction of 26S rRNA m^5^C methylation by 35% (Fig. 1C). In the NL2099 strain, *nsun-1* RNAi treatment also led to a reduction of 26S rRNA m^5^C methylation by 26% (Fig. 1D).

As there are only two known modified m^5^C residues on worm 26S rRNA, and since one of them is introduced by NSUN-5 (Adamla et al., 2019; Schosserer et al., 2015), a complete loss of NSUN-1 activity was expected to result in a 50% decrease in m^5^C methylation. However, protein depletion achieved with RNAi is usually not complete. It is not clear why the level of m^5^C depletion was not higher in the RNAi hypersensitive strain in comparison to the N2 strain; nonetheless, RNAi-mediated depletion of *nsun-1* significantly reduced the levels of 26S rRNA m^5^C modification in both worm strains, thus, we conclude that NSUN-1 is responsible for 26S rRNA m^5^C methylation.

We analysed the 26S rRNA m^5^C levels in a *nsun-5* deletion strain as control (strain JGG1, Fig 1E). In this case, we observed a near 2-fold reduction in methylation (58% residual), as expected from the known involvement of NSUN-5 in modification at position C2381. When *nsun-1* was additionally depleted by RNAi in the *nsun-5* knockout animals, the level of 26S rRNA m^5^C was further reduced to 43%, again in agreement with our conclusion that NSUN-1 is responsible for methylating the second position, C2982.

To further prove that NSUN-1 is not involved in C2381 modification, methylation levels at this position were specifically tested by Combined Bisulfite Restriction Analysis (COBRA) assay in animals depleted of *nsun-1* or *nsun-5*. This method is based on bisulfite conversion of total RNA, followed by PCR amplification and restriction digest, yielding two bands in case of methylation at C2381 and three bands in case of non-methylation (Adamla et al., 2019). As expected, only *nsun-5* depletion strongly reduced methylation at C2381, and there was no residual m^5^C2381 in the *nsun-5* knockout strain, while *nsun-1* RNAi had no effect on modification at this position (Fig. 1F). Bisulfite sequencing is well-known to be sensitive to RNA secondary structure (Warnecke et al., 2002), which likely explains why, despite repeated attempts, we could not monitor modification at position C2982 by use of this technique.

In conclusion, NSUN-1 and NSUN-5 are each responsible for installing one m^5^C onto the worm 26S rRNA, with NSUN-1 being responsible for position C2982 and NSUN-5 for position C2381 under the *bona fide* assumption that indeed only two m^5^C positions are present as described (Sharma and Lafontaine, 2015).

### The somatic tissue-specific depletion *of nsun-1* extends healthy lifespan

Next, we investigated if knockdown of *nsun-1* modulates healthy lifespan in a similar fashion as *nsun-5* (Schosserer et al., 2015). For this aim, we depleted *nsun-1* by RNAi in N2 wildtype animals starting from day 0 of adulthood and, surprisingly, did not observe any extension of mean or maximum lifespan (Fig. 2A). Consequently, we also evaluated the stress resistance of adult worms upon *nsun-1* or *nsun-5* depletion, as an increased health at an advanced age often improves resilience to adverse events (Lithgow et al., 1994). Indeed, depletion of either *nsun-1* or *nsun-5* increased resistance to heat stress compared to the RNAi control (Fig. 2B). Similarly, we tracked the movement of animals exposed to either empty vector control or RNAi directed against *nsun-1* or *nsun-5* in a time course analysis, starting at day 1 of adulthood up to day 16. Interestingly, we observed a trend towards increased average speed in all three independent experiments at day 8 of adulthood in both *nsun-1* (+47.8%) and *nsun-5* (+34.7%) depleted animals compared to the control, and, to a lesser extent at day 12 (*nsun-1* RNAi: +10.2%, *nsun-5* RNAi: +73.5% compared to the control) (Fig. 2C). Thus, while *nsun-1* knockdown does not extend lifespan, it improves the healthspan parameters thermotolerance and locomotion (Bansal et al., 2015; Rollins et al., 2017).

**Figure 2:**
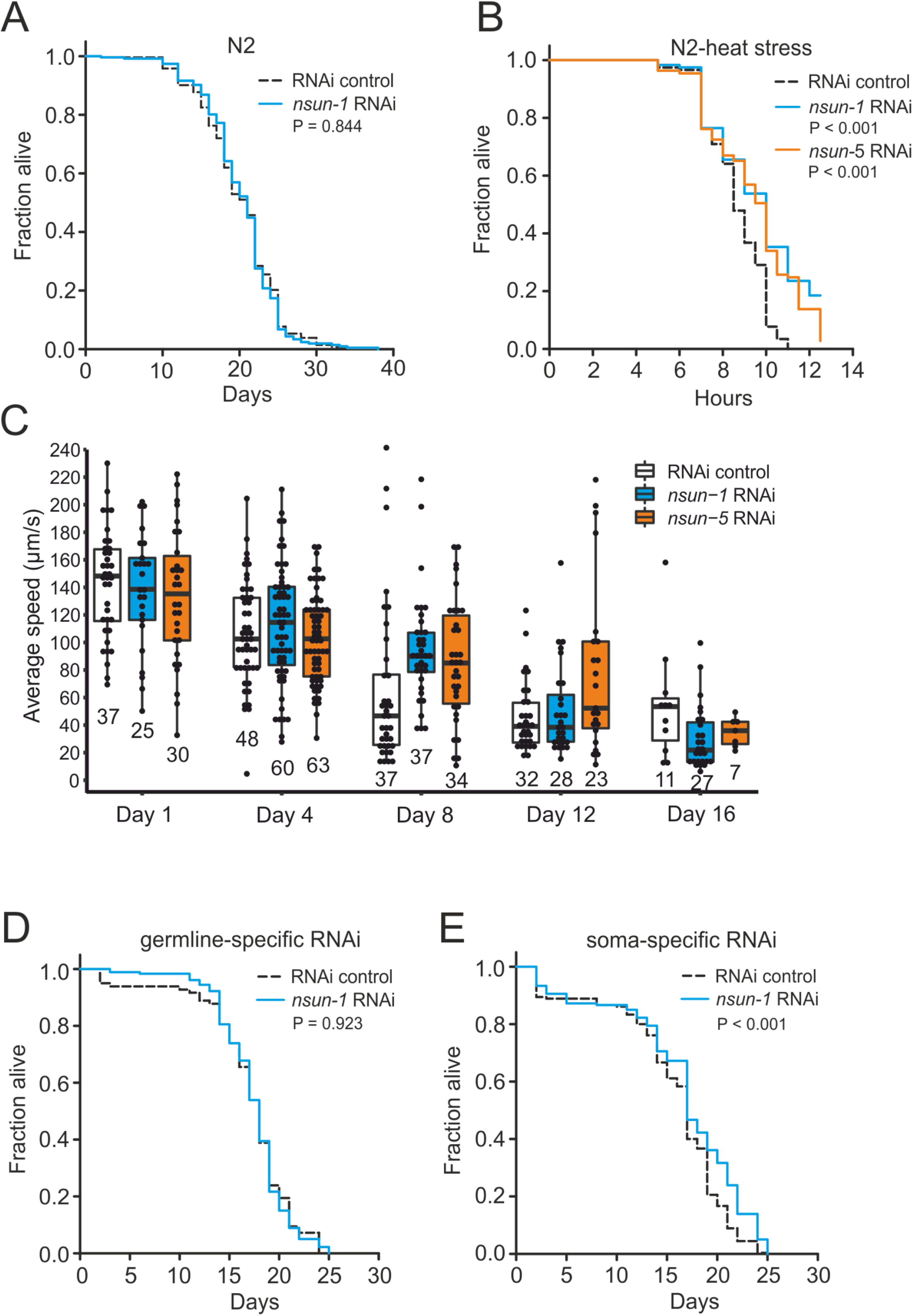
Depletion of *nsun-1* and *nsun-5* improves thermotolerance and locomotion. **A,** *nsun-1* whole-body RNAi (N2 wildtype strain) does not affect lifespan. Three pooled independent biological replicates are shown. Pooled n ≥ 208 animals per condition, log-rank test, not significant. **B,** N2 wildtype animals treated with either *nsun-1* or *nsun-5* RNAi and subjected to heat stress (35 °C) show increased survival compared to the RNAi control. Three pooled independent biological replicates are shown. Pooled n ≥ 90 animals per condition, log rank, P<0.05. **C,** Average speed [µm/s] of N2 wildtype worms as indicator of the health status was measured at day 1, 4, 8, 12 and 16 of adulthood. Movies of animals treated with either RNAi control, *nsun-1* or *nsun-5* RNAi were captured. One representative experiment is shown. Three biological replicates were performed with similar outcome. Pooled n ≥ 20 animals per condition at day 1. The black line indicates median. Two-way ANOVA P<0.01 (for interaction day:RNAi treatment) **D-E,** Lifespan analysis of germline- (NL2098) and soma-specific RNAi strains (NL2550). Worms were treated with either RNAi control or *nsun-1* RNAi. Only soma-specific knockdown of *nsun-1* results in increased lifespan **(E)** while germline-specific knockdown does not **(D)** (two independent biological replicates, pooled n(NL2098) ≥ 160 animals per condition, log-rank, not significant, pooled n(NL2550) ≥ 145 per condition, log-rank, P<0.01). A summary of the individual replicates of lifespan and thermotolerance experiments is provided in Figure 2–Source Data 1.

Intrigued that depletion of *nsun-1* did not extend the lifespan of *C. elegans* in a similar fashion as *nsun-5* did when whole adult animals were treated with RNAi, we reasoned that performing tissue-specific depletion of *nsun-1* might help us to further elucidate a possible effect on lifespan. We focused on the comparison of the germline and somatic tissues, because somatic maintenance and ageing are evolutionarily tightly connected (Kirkwood and Holliday, 1979), and signals from the germline modulate *C. elegans* lifespan (Hsin and Kenyon, 1999). In addition, only loss of soma- but not germline-specific eIF4E isoforms, which are central regulators of cap-dependent translation, extend nematode lifespan (Syntichaki et al., 2007). To test if *nsun-1* has similar specificity, we made use of worm strains sensitive to RNAi only in either the germline or somatic tissues. This is achieved, on the one hand by mutation of *rrf-1*, which is required for amplification of the dsRNA signal specifically in the somatic tissues (Kumsta and Hansen, 2012; Sijen et al., 2001), and, on the other hand by functional loss of the argonaute protein *ppw-1* rendering the germline resistant to RNAi (Tijsterman et al., 2002). Interestingly, germline-specific *nsun-1* RNAi had no effect on animal lifespan (Fig. 2D), but depletion of *nsun-1* in somatic tissues reproducibly increased mean lifespan by ∼10% (Fig. 2E).

In conclusion, both NSUN-1 and NSUN-5 m^5^C rRNA methyltransferases modulate thermotolerance and mobility of wildtype nematodes at midlife. Whole-animal *nsun-5* depletion expands mean lifespan by 17% (Schosserer et al., 2015), and in contrast to this, a 10% lifespan extension is only detected after depletion of *nsun-1* specifically in the somatic tissues.

### The somatic tissue-specific depletion of *nsun-1* affects body size, fecundity, and gonad maturation

The ‘disposable soma theory’ of ageing generally posits that long-lived species exhibit impaired fecundity and reduced number of progeny. The underlying cause is that energy is invested in the maintenance of somatic tissues instead of rapid reproduction (Kirkwood and Holliday, 1979). In keeping with this theory, we expected that the absence of *nsun-1* in somatic tissues, which increased longevity, may reduce fecundity. Therefore, we first measured the brood size upon *nsun-1* and *nsun-5* depletion by RNAi. After reaching adulthood but before egg-laying, worms were transferred to individual wells of cell culture plates containing NGM-agar and fed with bacteria expressing the specific RNAi or, as control, the empty vector. Egg production was impaired upon *nsun-1* knockdown (reduced by 42%), but, surprisingly, this was not the case upon *nsun-5* knockdown (reduced by 2%) (Fig. 3A). Egg production ceased rapidly after day one in all conditions (Figure 3–figure supplement 1A).

**Figure 3:**
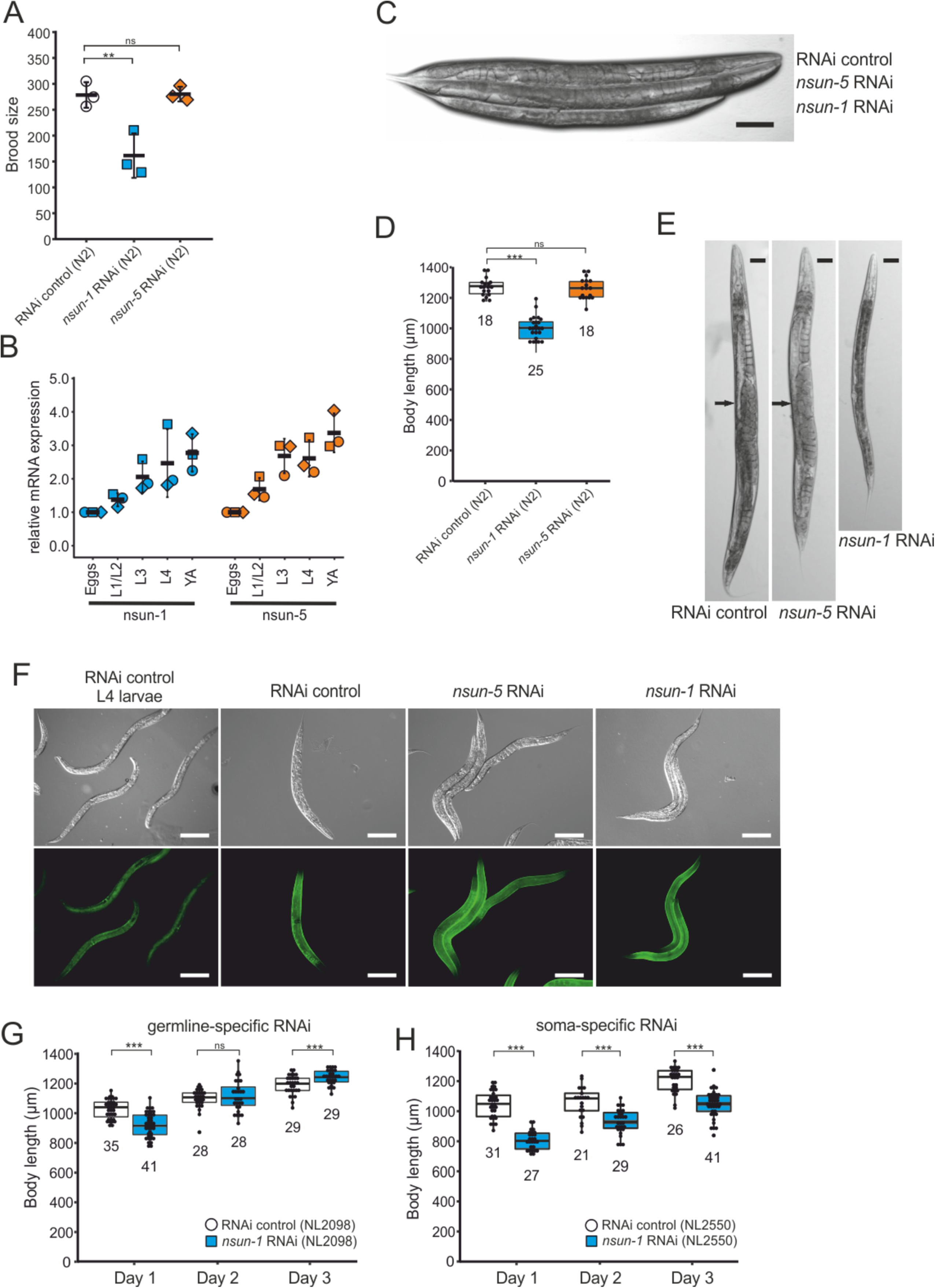
Loss of *nsun-1* reduces body size and impairs fecundity. **A,** Brood size analysis of adult-onset RNAi exposed animals. Eggs of individual worms were counted and the total number of eggs per worm is shown. Knockdown of *nsun-1* but not *nsun-5* induced a significant reduction in brood size compared to control RNAi (three independent experiments, n = 5 per condition and per experiment, one-way ANOVA with Dunnett′s post test, α=0.05, **P<0.01). Error bars indicate standard deviation. **B,** RT-qPCR analysis of wild-type animals at different stages of development (eggs, L1/L2 larvae, L3 larvae, L4 larvae and young adults). *tba-1* was used for normalization and expression is shown relative to eggs. Error bars represent standard deviation of three biological replicates, one-sample t-test against expected value of 1 with multiple comparison correction by Holm’s method did not reveal significant differences. **C,** Representative DIC images of larval-onset RNAi exposed nematodes show that only *nsun-1* but not *nsun-5* RNAi decreased the body length and altered general morphology compared to the RNAi control. Scale bar, 100 µm. **D,** Quantification of mean body length of 1-2-day old adult worms. The body size of *nsun-1* RNAi treated worms was significantly reduced compared to the RNAi control and *nsun-5* RNAi. The experiment was independently performed two times and one representative replicate is shown. n(RNAi control) = 18, n(*nsun-1* RNAi) = 25, n(*nsun-5* RNAi) = 19, one-way ANOVA with Dunnett’s post, α=0.05, ***P<0.001. Error bars represent standard deviation. **E,** Larval-onset *nsun-1* RNAi-treated adults had reduced body size and lacked embryos (arrow). Scale bar, 50 µm. **F,** Loss of *nsun-1* did not impair expression of the adult-specific marker col-19::GFP. The TP12 strain was used and young adult animals treated with either control RNAi, *nsun-1* RNAi or *nsun-5* RNAi were imaged in DIC and fluorescent mode. L4 control RNAi worms, which did not express GFP specifically in the hypodermis, were used as negative control. Scale bar, 200 µm. **G-H,** Soma- but not germline-specific *nsun-1* RNAi phenocopied the mean body length defect upon whole body *nsun-1* knockdown. The germline-specific NL2098 strain **(G)** and the soma specific NL2550 **(H)** strain were used and measured on three consecutive days after reaching adulthood. n ≥ 21 for each day and condition. Two independent experiments were performed and one representative replicate is shown. Two-tailed t-test, ***P<0.001. Error bars represent standard deviation.

Thus far, all the experiments were performed on worms subjected to adult-onset *nsun-1* knockdown, as animals depleted of *nsun-1* during development were smaller and were infertile upon adulthood. To follow up on these observations, we measured mRNA expression levels of both m^5^C rRNA methyltransferases at different developmental stages including eggs, L1/L2 larvae, L3 larvae, L4 larvae and young adults. RT-qPCR indicated that both *nsun-1* and *nsun-5* mRNA levels constantly increase during development (Fig. 3B). The same observation applied to mRNA levels of *nsun-2* and *nsun-4* (Figure 3–figure supplement 1B), indicating that all four members of the NSUN-protein family might play important roles during development.

To further assess whether *nsun-1* expression is indeed necessary for progressing faithfully through larval stages, we captured images of young adult animals subjected to larval-onset RNAi. The disparity in body size between RNAi control and *nsun-1* RNAi was apparent, whereby *nsun-1* depleted animals showed reduced length by approximately 20%. Interestingly, this reduced body size was not seen upon *nsun-5* RNAi treatment (Fig. 3C,D).

In addition, we imaged 3-day old animals using differential interference contrast (DIC) microscopy. Worms subjected to *nsun-1* knockdown displayed morphological alterations, specifically the gonad appeared severely distorted (Fig. 3E). In contrast, RNAi control and *nsun-5* RNAi showed comparable morphology of distal and proximal gonads (Fig. 3E). Consequently, we hypothesized that *nsun-1* RNAi treated worms might be arrested in early L4 larval stage when the gonad is not yet fully developed and animals still grow, instead of reaching normally adulthood after 3 days like RNAi control or *nsun-5* RNAi treated nematodes. To test this, we used the TP12 *kaIs12[col-19::GFP]* translational reporter strain, which expresses COL-19::GFP specifically upon reaching adulthood, but not during larval stages. Surprisingly, larval-onset RNAi against *nsun-1* or *nsun-5* did not reveal differences in the expression of COL-19::GFP as compared to RNAi control, suggesting that neither *nsun-1* nor *nsun-5* induce larval arrest (Fig. 3F). Together with the reduced brood size upon adult-onset RNAi, these findings imply that loss of *nsun-1* induces phenotypic changes in the reproductive organs of *C. elegans* independent of development.

Since knockdown of *nsun-1* extended lifespan only when it was applied to somatic tissues, we hypothesized that body length might also be affected when these tissues are specifically targeted for depletion. To test this, we depleted *nsun-1* specifically in the germline or in somatic tissues using tissue-specific RNAi strains and measured body size during three consecutive days after adulthood was reached. While germline-specific knockdown of *nsun-1* did not induce any changes in body size (Fig. 3G), soma-specific knockdown revealed a decrease in body length by 23% on day 1, 11% on day 2 and 12% on day 3 compared to the RNAi control (Fig. 3H), phenocopying *nsun-1* depletion in wildtype animals after whole animal RNAi. Similarly, no effect of *nsun-1* knockdown was evident in another germline-specific RNAi strain, which was recently developed to enhance germline specificity (Zou et al., 2019) (Figure 3–figure supplement 1C).

In conclusion, *nsun-1* but not *nsun-5* depletion impairs body size and morphology of the gonad and leads to a significant reduction of brood size. Furthermore, these phenotypes are also observed when *nsun-1* is specifically knocked-down in somatic tissues, but not when depleted in the germline only.

### NSUN-1 is required for the transition of meiotic germ cells to mature oocytes

To further investigate the mechanisms underlying impaired fecundity upon *nsun-1* knockdown, we analysed the morphology of the gonad in *nsun-1* depleted animals in more detail. The germline of adult hermaphrodites resides within the two U-shaped arms of the gonad, which contains germ cells at various stages of differentiation (Fig. 4A). The gonad is sequentially developing from the proliferative germ cells near the distal tip cell, through the meiotic zone into the loop region, finally culminating in fully-formed oocytes in the proximal gonad (Pazdernik and Schedl, 2013). The limiting factor for fecundity in self-fertilizing hermaphrodites is sperm produced in the spermatheca (Hodgkin and Barnes, 1991).

**Figure 4:**
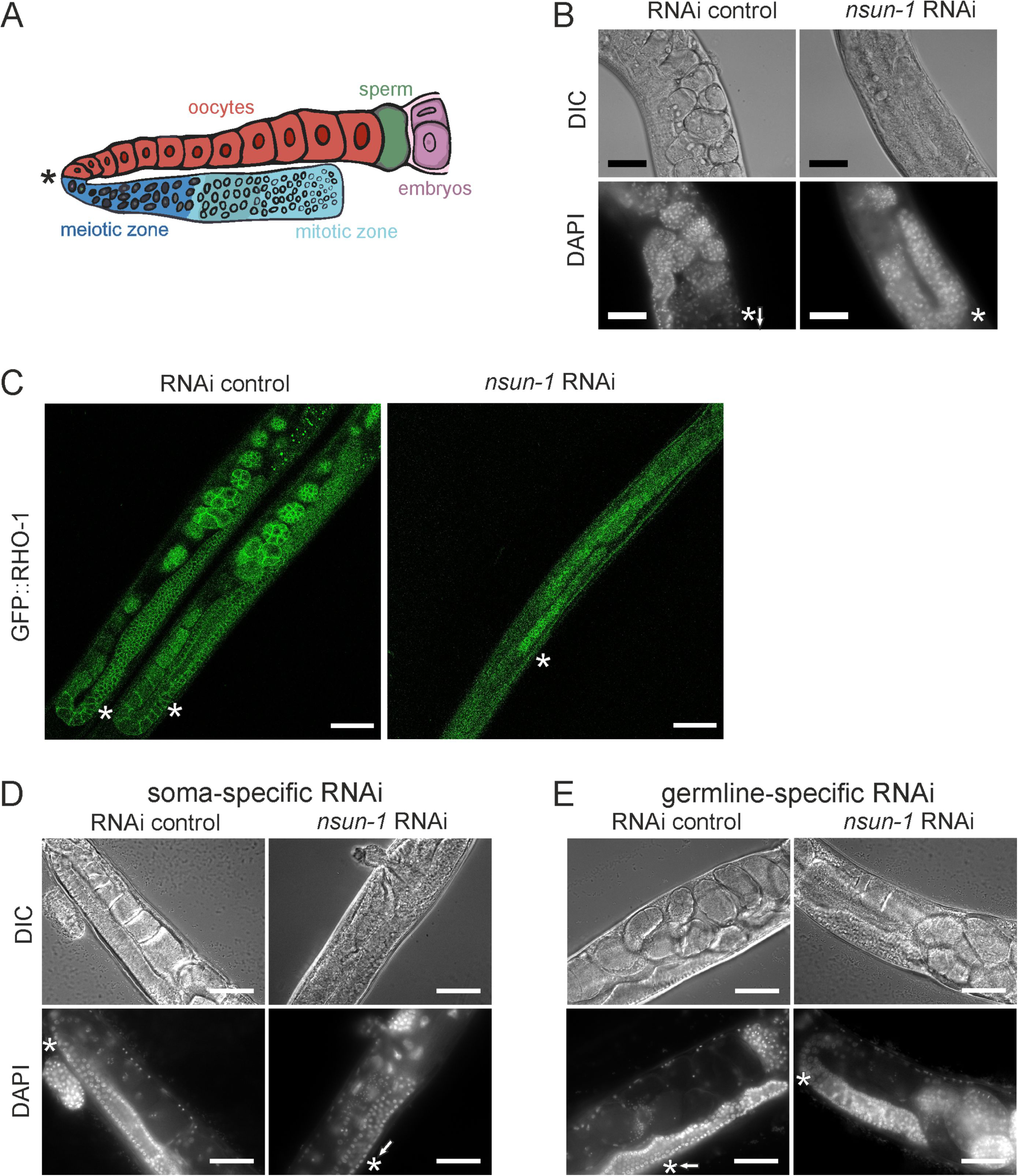
Soma-specific depletion of *nsun-1* blocks oogenesis. **A,** Schematic of one gonad arm in *C. elegans*. Germ cell replication starts in the distal mitotic zone. After passing through the meiotic zone, oocytes further mature and are fertilized by sperm produced in the spermatheca. In panels **A-E**, an asterisk indicates the gonadal region impaired in *nsun-1* RNAi exposed animals. This area corresponds to the transition between the meiotic zone and oocyte maturation. **B,** Microscopic image of one gonad arm of young adult worms subjected to either control or *nsun-1* RNAi. Worms were imaged in DIC mode and nuclei of fixed animals were stained with DAPI. Scale bar, 40 µm. **C,** Confocal imaging of the gonad-specific GFP::RHO-1 expressing SA115 strain revealed altered gonad morphology upon *nsun-1* knockdown (see also Figure 4–figure supplement 1). Scale bar, 40 µm. **D-E,** Soma- but not germline-specific *nsun-1* RNAi phenocopied altered gonad morphology upon whole body *nsun-1* depletion. NL2550 was used for soma- **(D)** and NL2098 for germline-specific knockdown **(E)**. One gonad arm of one representative 2-day old adult animal was imaged in DIC mode and nuclei were stained with DAPI following fixation. Scale bar, 40 µm.

Upon visualizing the germline cell nuclei with DAPI-staining, no oocytes were observed in worms after knockdown of *nsun-1* in contrast to RNAi control treated animals (Fig. 4B). The mitotic zone at the distal end of the gonad appeared normal in *nsun-1* depleted animals, whereas oocyte production starting at the pachytene zone was hampered. Analysis of GFP::RHO-1 and NMY-2:GFP expressing worm strains, which specifically express GFP in the germline, confirmed our observations (Fig. 4C, Figure 4–figure supplement 1). The gonads of control and *nsun-5* RNAi treated animals appeared normal, clearly depicting the different stages of *in-utero* embryo development, whereas the germline of *nsun-1* RNAi treated animals displayed a strikingly altered morphology.

Since other phenotypes observed upon *nsun-1* depletion were detected in soma- but absent from germline-specific RNAi treated strains, we hypothesized that the somatic part of the gonad might specifically require NSUN-1 for normal oocyte production. Indeed, soma-specific depletion of *nsun-1* phenocopied the distorted gonad morphology of wildtype animals exposed to *nsun-1* RNAi (Fig. 4D). Remarkably, upon germline-specific knockdown of *nsun-1,* the gonad appeared completely unaffected (Fig. 4E; Figure 4–figure supplement 2).

### NSUN-1 is not essential for pre-rRNA processing and global protein synthesis

Since the only known function of NSUN-1 and NSUN-5 is m^5^C methylation of rRNA, we reasoned that methylation-induced alterations of ribosome biogenesis and function might explain the observed phenotypes. Therefore, we tested if the presence of NSUN-1 or NSUN-5 is required for ribosomal subunit production and pre-rRNA processing. To this end, total RNA was extracted from worms treated with *nsun-1* RNAi, separated by denaturing agarose gel electrophoresis and processed for northern blot analysis (Fig. 5A,B). Again, two reference worm strains were used (N2 and NL2099). Upon *nsun-1* knockdown, we observed a mild accumulation of the primary pre-rRNA transcript and of its immediate derivative, collectively referred to as species “a” (Fig. 5A, (Bar et al., 2016; Saijou et al., 2004)), as well as a mild accumulation of the pre-rRNAs “b” and “c’” (Fig. 5A,B, see lane 1 and 3 as well as 5 and 6). Again, these findings were observed in both worm backgrounds tested, N2 and NL2099.

**Figure 5:**
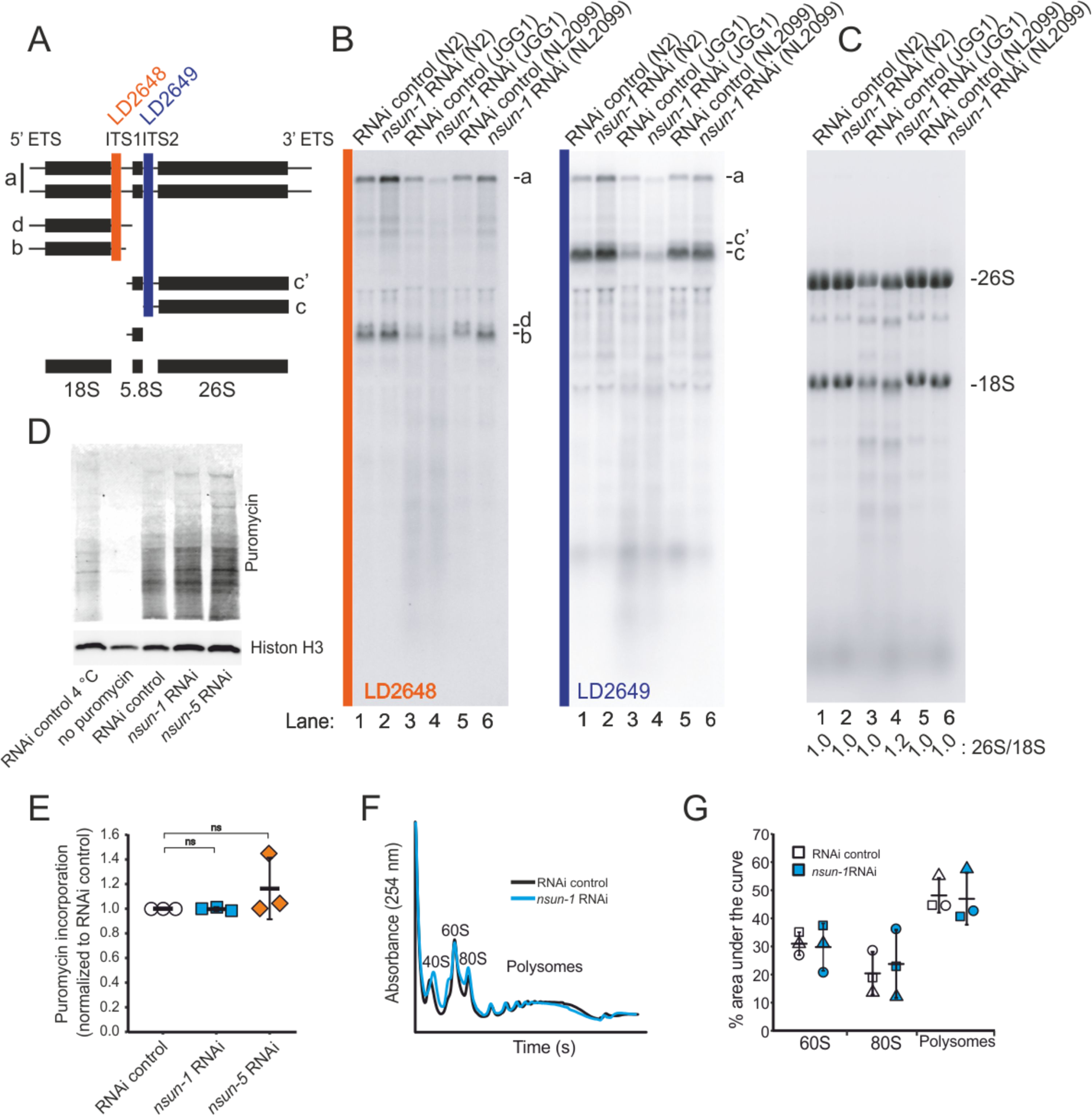
NSUN-1 and NSUN-5 are only partially required for rRNA processing and not for global translation. **A,** Schematics of pre-rRNA processing intermediates in *C. elegans* and probes (LD2648 and LD2649) used in pre-rRNA processing analysis (see panel B). **B,** Pre-rRNA processing analysis. Total RNA extracted from the indicated strains were separated on denaturing agarose gels and processed for northern blotting. The probes (LD2648 and LD2649) used to detect the pre-rRNA intermediates a, b, c, ć, and d are indicated. **C,** Steady-state levels of mature rRNAs (18S and 26S) analysed by ethidium bromide staining and quantified by densitometry. The 26S/18S ratio is indicated. **D,** Total protein synthesis of N2 animals treated with RNAi control, *nsun-1* or *nsun-5* RNAi. RNAi control treated worms at either 4°C or without puromycin exposure were used as negative controls. Protein synthesis was measured by puromycin exposure for 3 h and western blot using a puromycin-specific antibody. The experiment was performed in three independent replicates. One representative replicate is shown. Histone H3 was used as loading control. **E,** Quantification of western blots in D (three biological replicates, one-sample t-test against an expected value of 1, α=0.05, not significant). **F-G,** Polysome analysis indicating that global translation is not affected by *nsun-1* depletion. Free small subunit (40S), large subunit (60S), monosome (80S) and polysome fractions were detected by UV_254_ monitoring. Representative profiles are shown. **G,** Quantification of 60S, 80S and polysome fractions of three independent experiments reveals no changes between *nsun-1* knockdown and RNAi control.

For comparison, we also analysed rRNA processing in *nsun-5* deletion worms (JGG1 strain) in presence and absence of NSUN-1 (*nsun-1* RNAi in JGG1). In both cases, we noted an important reduction in the overall production of ribosomal RNAs (Fig. 5B,C), with an apparent increase of rRNA degradation (seen as an increase in accumulation of metastable RNA fragments, in particular underneath the 18S rRNA). Furthermore, we observed that NSUN-1 is not required for mature rRNA production as shown by the unaffected levels of mature 18S and 26S rRNAs (Fig. 5C). This was confirmed by determining the 26S/18S ratio, which was 1.0 as expected since both rRNAs are produced from a single polycistronic transcript (Fig. 5C). The levels of the other two mature rRNAs (5S and 5.8S) were also unaffected (Figure 5–figure supplement 1). This behaviour was shown in both worm backgrounds, N2 and NL2099, used. The overall decrease in mature ribosome production observed in *nsun-5* deletion worms did not affect the ratio of mature ribosomal subunits (26S/18S ratio of 1.0) (Fig. 5C). In agreement with the reduced amounts of 18S and 26S rRNA observed in *nsun-5* deletion worms, total amounts of all precursors detected were reduced (Fig. 5B). Analysis of low molecular weight RNAs by acrylamide gel electrophoresis, revealed absence of NSUN-5 to severely inhibit processing in the internal transcribed spacer 2 (ITS2), which separates the 5.8S and 26S rRNAs on large precursors. This was illustrated by the accumulation of 3’-extended forms of 5.8S, and of short RNA degradation products (Figure 5–figure supplement 1, see lanes 3 and 4). Depletion of *nsun-1* partially suppressed the effect of *nsun-5* deletion: the overall production of mature rRNA and, in particular, the amount of mature 26S rRNA was increased (ratio of 1.2) (Fig. 5C). Consistently, the accumulation of 3’-extended forms of 5.8S and of short RNA degradation products was reduced (Figure 5–figure supplement 1).

In order to test if mature ribosomes of animals lacking any of the two m^5^C rRNA methyltransferases might be functionally defective, we analysed global protein synthesis by incorporation of puromycin in N2 worms treated with either RNAi control, *nsun-1* or *nsun-5* RNAi. Worms were exposed to puromycin for three hours at room temperature. Following lysis, puromycin incorporation was measured by western blot with an anti-puromycin antibody (Fig. 5D). Quantification of three independent experiments revealed no changes in global protein synthesis (Fig. 5E). We also performed polysome profiling which provides a “snapshot” of the pool of translationally active ribosomes. Comparison and quantification of profiles obtained from control and *nsun-1* knockdown nematodes did not reveal any differences in the distribution of free subunits, monosomes and polysomes (Fig. 5F,G). This agrees with the absence of global protein translation inhibition in the metabolic (puromycin) labelling assay (Fig. 5E).

In conclusion, NSUN-1 is not required for rRNA processing nor for global translation. On the contrary, the amounts of ribosomal subunits were reduced in the absence of NSUN-5. The ribosomal biogenesis alterations observed upon *nsun-5* depletion result from a combination of processing inhibitions in ITS2 and increased rRNA intermediates turnover. However, global translation was not detectably affected.

### mRNAs encoding cuticle collagens are translationally repressed upon *nsun-1* **knockdown**

Since depletion of *nsun-1* did not affect global protein synthesis, we hypothesized that loss of 26S rRNA m^5^C methylation might modulate the translation of specific mRNAs, as was previously observed after Rcm1 (NSUN-5 homolog) depletion in yeast (Schosserer et al., 2015). To test this possibility, we isolated mRNAs contained in the polysome fraction, systematically sequenced them by RNAseq and compared their respective abundance in polysomes versus total mRNAs contained in the lysate before fractionation. We considered only protein-coding mRNAs with a minimum fold change of 2 between translatome and transcriptome and an adjusted p-value cut-off at 0.05 (Fig. 6A; Supplemental Data File 1). The translation of more mRNAs was repressed (RNAi control: 599, *nsun-1* RNAi: 536) than promoted (RNAi control: 94, *nsun-1* RNAi: 84). Since the composition of 3’ UTRs can affect translation (Tushev et al., 2018), we analysed GC-content, length and minimal free folding energy of all coding, promoted and repressed mRNAs in our dataset (Fig. 6B). Interestingly, all three features significantly differed between RNAi control and *nsun-1* RNAi in promoted and repressed mRNAs (p < 0.05), while they remained unchanged when analysing all coding mRNAs present in the dataset. These findings suggest that loss of *nsun-1* causes the translation of specific subsets of mRNAs based on the composition and length of their 3’UTRs. Moreover, 3’UTRs of mRNAs repressed by *nsun-1* depletion were exclusively and significantly enriched (p < 0.001) for several binding motifs of ASD-2, GLD-1 and RSP-3 (Supplemental Data File 2). All three RNA binding proteins play essential roles in *C. elegans* development (Lee and Schedl, 2010; Longman et al., 2000) and were not differentially regulated in the transcriptome between *nol-1* RNAi and RNAi control (Supplementary Data File 3).

**Figure 6:**
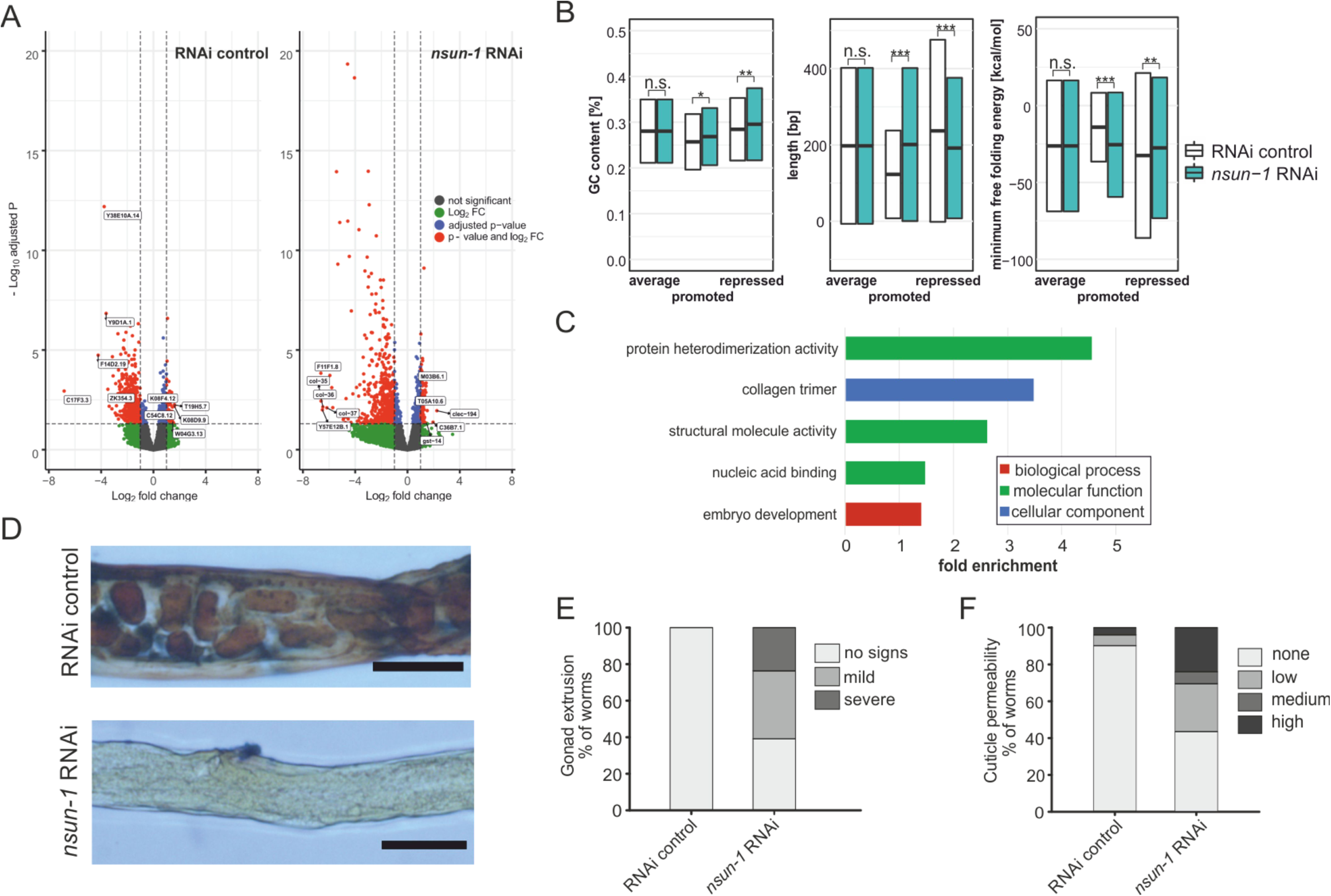
*nsun-1* depletion modulates selective translation of collagens and induces gonad extrusion and loss of barrier function. **A,** Vulcano plots of selectively translated genes after RNAi control and *nsun-1* RNAi exposure. Significantly regulated genes (adjusted p < 0.05 and fold-change > 2) between polysome fraction and total mRNAs are depicted in red, genes with a two-fold up- or down-regulation but an adjusted p-value (FDR/ Benjamini and Hochberg) above 0.05 in green, genes with an adjusted p-value below 0.05 but less than two-fold change in expression in blue, and not significantly regulated genes in grey. The top five up- or down-regulated genes based on their fold change are indicated. **B,** Characteristics of the 3’UTRs of mRNAs with significantly promoted or repressed polysome enrichment (adjusted p < 0.1, fold-change > 2). GC content (in %), length (in bp) and minimum free folding energy [kcal/mol] are shown. Boxes indicate mean ± SD (for GC content and length) or mean ± SEM (for minimum free folding energy). Wilcoxon rank sum test, *P<0.05, **P<0.01, ***P<0.001. **C,** Biological GO terms enriched among genes with repressed translation (adjusted p < 0.1, fold-change > 2) upon *nsun-1* depletion. Modified Fisher’s exact test, p < 0.05. **D,** Histological staining (Herovici) to assess collagen deposition. Worms exposed to control RNAi show presence of both young (blue) and mature (pink to brownish-red) collagen whereas animals subjected to *nsun-1* RNAi display less collagen deposition. The cytoplasm is counterstained in yellow. Representative images of the region surrounding the gonad are shown. Two independent experiments with a minimum of 10 animals each were performed with similar outcome. Scale-bar, 80 µm. **E,** Quantification of gonad extrusion upon *nsun-1* depletion compared to RNAi control. 8-9 day old adult animals were classified into three categories according to the severance of gonad extrusion (‘no signs’, ‘mild’, ‘severe’, see Figure 6–figure supplement 1A). The experiment was independently performed two times with similar outcome. One representative replicate is shown. n ≥ 50 animals per replicate. Modified Fisher’s exact test on the raw count values, p < 0.001. **F,** Quantification of cuticle barrier function upon *nsun-1* depletion compared to RNAi control. Young adult animals were exposed to Hoechst 33342, which is membrane-permeable but cuticle-impermeable. Stained nuclei were counted exclusively in the tail region to exclude intestinal autofluorescence and classified into four categories accordingly (‘none’, ‘low’, ‘medium’, ‘high’, see Figure 6–figure supplement 1B). Three independent experiments were pooled. n(RNAi control) = 51, n(*nsun-1* RNAi) = 46. Modified Fisher’s exact test on the raw count values, p < 0.001.

To further understand the mechanistic link between differential translation and the phenotypes observed upon *nsun-1* knockdown, we performed GO-term enrichment analysis. Among others, GO terms associated with collagens, structural integrity of the cuticle and embryo development were significantly enriched amongst those mRNAs, which were translationally repressed by *nsun-1* knockdown (Fig. 6C, Supplemental Data File 4).

Since three collagens (*col-35*, *col-36* and *col-37*) were also among the five most strongly repressed genes upon loss of *nsun-1* (Fig. 6A), we decided to assess whether collagen deposition is indeed altered. For this aim, we performed a specific histological staining protocol in which young collagen is strained blue and mature collagen is stained pink to brownish-red (Herovici, 1963; Teuscher et al., 2019). While young adult worms exposed to RNAi control showed presence of both young and mature collagen, animals subjected to *nsun-1* RNAi displayed a strikingly reduced collagen deposition compared to the cytoplasmatic counter-stain (yellow) (Fig. 6D).

Interestingly, we repeatedly observed an increased fraction of animals displaying gonad extrusion upon *nsun-1* RNAi, which might be caused by loss of cuticle structural integrity. To quantify this phenotype, we classified mid-aged animals according to the grade of gonad extrusion into three categories: i) no visible signs of gonad extrusion, ii) mild extrusion, or iii) severe extrusion (Figure 6–figure supplement 1A). Upon *nsun-1* depletion, 134 of 220 animals (∼60%) showed mild to severe extrusion of the gonad (categories ii and iii), while no extrusion was observed in any of the tested 50 RNAi control nematodes (Fig. 6E). To assess further physiological consequences of altered collagen deposition, we tested cuticle barrier activity. This assay is based on the principle Hochst33342 dye being membrane permeable, but cuticle impermeable. As previously described by Ewald et al. 2015, worms were grouped into four categories: i) not permeable (no stained nuclei in the animal tail region), ii) mildly permeable (< 5 stained nuclei), iii) permeable (5-10 stained nuclei), or iv) highly permeable (>10 stained nuclei) (Figure 6–figure supplement 1B,C). Consistent with a change of several collagens, *nsun-1* RNAi caused cuticle permeability (categories ii, iii and iv) in 26 of 46 animals (∼56%) compared to only 5 of 51 RNAi control animals (∼10%) (Fig. 6F).

Taken together, this indicates that NSUN-1 is partially required for translation of several cuticle collagens, which might not only explain the loss of gonad integrity, but also the increased cuticle permeability upon *nsun-1* depletion. Moreover, several mRNAs whose translation depends on NSUN-1 are associated with embryogenesis and enriched for binding motifs of known regulators of nematode development.

## Discussion

Although rRNAs are universally modified at functionally relevant positions, little is known about the biological functions and pathological roles of RNA methylation sites, or about their potential readers, writers and erasers (Sharma and Lafontaine, 2015). In this work we have investigated the molecular and physiological roles of two structurally related Sun-domain-containing RNA methyltransferases, NSUN-1 and NSUN-5 in *C. elegans*, each responsible for writing one m^5^C mark on 26S rRNA. We further describe NSUN-1 as a *bona fide* m^5^C rRNA writer enzyme that, if missing, directly entails physiological and developmental consequences. We conclude that, molecularly, loss of NSUN-1 function leads to translational remodelling with profound consequences on cell homeostasis, exemplified by loss of cuticle barrier function, and highly specific developmental defects, including oocyte maturation failure. We further suggest that extrusion of the gonad and loss of cuticle barrier function are directly caused by reduced expression of collagens, while the developmental defects are associated with altered expression of several important developmental regulators. We summarized the observed RNAi phenotypes in different strain backgrounds in Table 1.

**Table 1:** Comparison of phenotypes after nsun-1 and nsun-5 depletion (**in green**) soma-specific effects, n.d.: not determined.

| Phenotype | *nsun-1* RNAi | *nsun-5* RNAi |
| --- | --- | --- |
| Lifespan | - Unaffected in whole adult treatment<br>- Unaffected after germline-specific depletion<br>- Increased by ~10% after soma-specific depletion | - Increased by ~17% in whole adult treatment (Schosserer et al., 2015) |
| Stress resistance (heat) in adults | Increased | Similarly increased |
| Locomotion at midlife | Increased | Similarly increased |
| Brood size (fecundity) | Reduced (2-fold) | Unaffected |
| Adult animal size | - Reduced by ~20% after soma-specific depletion and whole body depletion<br>- Unaffected after germline-specific depletion | Unaffected |
| Gonad morphology | - Impaired at meiotic to oocyte transition<br>-Gonad extrusion (possibly caused by loss of cuticle integrity)<br>- Unaffected after germline- | Unaffected |
|  | specific depletion |  |
| Pre-rRNA processing | Unaffected | Affected |
| Collagen expression | Affected (translational remodeling) | n.d. |
| Cuticle permeability | increased | n.d. |

According to the ‘disposable soma theory of aging’, a balance between somatic repair and reproduction exists. Depending on the current environment, an organism may direct the available energy either to maintenance of the germline and thereby ensuring efficient reproduction, or to the homeostasis of somatic cells including the prevention of DNA damage accumulation (Kirkwood and Holliday, 1979). While most of the known genetic and nutritional interventions increase the lifespan of organisms, they antagonistically also reduce growth, fecundity and body size (Kapahi, 2010; Kenyon et al., 1993). Indeed, reduction of overall protein synthesis by genetic, pharmacological or dietary interventions was reproducibly shown to extend longevity in different ageing model organisms (Chiocchetti et al., 2007; Curran and Ruvkun, 2007; Hansen et al., 2007; Kaeberlein et al., 2005; Masoro, 2005; Pan et al., 2007). These reports clearly established protein synthesis as an important regulator of the ageing process at the interface between somatic maintenance and reproduction. Thus, we were surprised to find that despite their ability to modulate healthy ageing and to methylate rRNA, neither NSUN-1 nor NSUN-5 were required for global protein synthesis in worms under the conditions tested. In the case of NSUN-5 depletion, we previously found overall translation to be decreased in mammalian cells (Heissenberger et al., 2019), but not in yeast (Schosserer et al., 2015). We reasoned that the higher complexity of mammalian ribosomes and associated factors might render them more vulnerable to alterations of rRNA secondary structure, for example by loss of a single base modification, than ribosomes from yeast and nematodes.

As ribosome biogenesis or global translational activity *per se* were not severely affected by loss of NSUN-1, the idea of a mechanism to ‘specialize’ the ribosome by rRNA modifications for selective mRNA translation seems attractive (Simsek and Barna, 2017). Indeed, lack of NSUN-1 and presumably the methylation at C2982 resulted in decreased translation of mRNAs containing GLD-1 and ASD-2 binding sites. These two proteins are closely related members of the STAR protein family involved in mRNA binding, splicing and nuclear export of mRNAs. While the molecular functions of ASD-2 are only poorly understood, the role of GLD-1 in embryonic development is well characterized (Lee and Schedl, 2010). GLD-1 levels are highest in the pachytene (also referred to as meiotic zone), where GLD-1 acts as a translational repressor of mRNAs modulating oogenesis. At the transition zone between the pachytene and the diplotene, GLD-1 levels sharply decrease and previously repressed mRNAs are consequently translated. Since the gonads of *nsun-1* knockdown animals appeared defective precisely at this transition and GLD-1 target mRNAs were repressed, we speculate that either ribosomes lacking the methylation at C2982 have generally low affinity for these mRNAs, or that translational repression by GLD-1 is never fully relieved. Although the expression of GLD-1 itself was not differentially regulated between control and *nsun-1* depleted animals at transcription or translation level (Supplemental Data File 1 and 3), multiple direct or indirect interactions with NSUN-1 to modulate ribosome function are still conceivable and will require further studies.

Previously, Curran and Ruvkun reported that depletion of *nsun-1* (W07E6.1) by adult-onset RNAi was able to extend lifespan in *C. elegans* (Curran and Ruvkun, 2007). The authors used for their high throughput screen a strain carrying a mutation in the *eri-1* gene, rendering it hypersensitive to RNAi in the whole body, but especially in neurons and in the somatic gonad (Kennedy et al., 2004). In this study, we conducted a whole-body knockdown in N2 wildtype animals, but could not verify these previous findings on lifespan, although thermotolerance and the health status of mid-aged nematodes, as assessed by quantifying locomotion behaviour, were elevated. However, when knocking-down *nsun-1* specifically in somatic tissues, but not in the germline, we observed a lifespan extension. The N2 wildtype strain is usually resistant to RNAi in the somatic gonad and neurons. Thus, we hypothesize that depletion of *nsun-1* specifically in the somatic part of the gonad is required for lifespan extension, which is only effectively realised in the *eri-1* and *ppw-1* mutant strains, but not in N2 wildtype animals. Intriguingly, these findings further suggest possible non-cell-autonomous effects of single RNA methylations, since modulation of NSUN-1 levels in somatic cells profoundly affected distinct cells of the germline.

The developing gonad of L1 larvae consists of two primordial germ cells and two surrounding somatic gonad precursor niche cells. The crosstalk between these two cell types, which form the germline and somatic part of the gonad at later stages of larval development, was already described to modulate ageing and stress responses. Laser depletion of both primordial germ cells extends lifespan via insulin/IGF-signalling, while animals with an additional depletion of the two somatic gonad precursor cells have a normal lifespan (Hsin and Kenyon, 1999). Of potential relevance to our study is a recent report by Ou and coworkers, who demonstrated that IFE-4 regulates the response to DNA damage in primordial germ cells in a non-cell-autonomous manner via FGF-like signalling. Soma-specific IFE-4 is involved in the specific translation of a subset of mRNA including *egl-15*. Thereby, IFE-4 regulates the activity of CEP-1/p53 in primordial germ cells despite not being present there (Ou et al., 2019). We thus hypothesize that selective translation of mRNAs by specialized ribosomes, either generated by association with translational regulators such as IFE-4, or by RNA modifications as described here, might serve as a general mechanism to tightly control essential cellular processes even in distinct cells and tissues.

Elucidating the precise mechanism is of importance, as human NSUN1 (also known as NOP2 or P120) was shown to be required for mammalian preimplantation development (Cui et al., 2016) and thus indicates evolutionary conservation. Cui and colleagues found that NSUN1 has an indispensable role during blastocyst development within their experimental setup. Additionally, other groups reported that low levels of NSUN1 reduce cell growth in leukaemia cells, which is in line with findings that NSUN1 promotes cell proliferation. Moreover, high NSUN1 expression results in increased tumour aggressiveness and augmented 5-azacytidine (5-AZA) resistance in two leukaemia cell lines (Bantis et al., 2004; Cheng et al., 2018; Saijo et al., 2001). Thus, NSUN1 might be considered as an example of ‘antagonistic pleiotropy’. According to this theory, genes can be indispensable early in life but negligible later, for instance after sexual reproduction. While NSUN1 appears to be essential for normal development, it might increase tumour aggressiveness later in life, especially in highly proliferative cells and tissues.

## Material and Methods

### Worm strains and culture conditions

The following *C. elegans* strains were used in this study: N2; JGG1 *nsun-5(tm3898)*, SA115 *unc-119(ed3)*, JJ1473 *unc-119(ed3)*, TP12 *kaIs12[col-19::GFP]*, DCL569 *mkcSi13[P_sun-1_::rde-1::sun-1 3’UTR + unc-119(+)]*, NL2098 *rrf-1(pk1417)* and NL2550 *ppw-1(pk2505)*. Worms were cultured following standard protocols on *Escherichia coli* OP50-seeded NGM agar plates at 20°C, unless indicated otherwise (Brenner, 1974).

### RNAi knockdown

The *nsun-1* RNAi clones was from the J. Ahringer library (Dong et al., 2003) and the *nsun-5* RNAi clone from the M. Vidal library (Rual et al., 2004). For inactivating *nsun-1* and *nsun-5*, feeding of double-stranded RNA expressed in bacteria was used (Timmons et al; 2001). Therefore, the HT115 strain of *E. coli,* carrying either the respective RNAi construct or the empty vector (L4440) as RNAi control, was cultured overnight in LB medium with ampicillin and tetracyclin at 37°C. Bacteria were harvested by centrifugation, resuspended in LB medium and either 100 µL (60 mm plates) or 400 µL (100 mm plates) were plated on NGM containing 1 mM isopropyl-b-D-thiogalactoside and 25 µg/mL carbenicillin. The plates were incubated at 37°C overnight and used within one week.

Larval-onset RNAi was achieved by bleaching adult animals. Released eggs were transferred directly to plates seeded with RNAi bacteria. Adulthood was usually reached after three days and animals were used for experiments when the RNAi control strain started to lay eggs.

In case of adult-onset RNAi, eggs were transferred to plates seeded with RNAi control bacteria. Animals were raised until egg production commenced and subsequently transferred to the respective RNAi bacteria.

### Differential Interference Contrast (DIC) microscopy

Worms were paralyzed using 1 M sodium azide solution and mounted on 2% agar pads. Images were acquired on a Leica DMI6000B microscope with a 10x dry objective (NA 0.3) or a 63x glycerol objective (NA 1.3) in DIC brightfield mode. Cropping, insertion of scale bars and brightness and contrast adjustments were done with Image J (version 2.0.0-rc-65/1.51w; Java 1.8.0_162 [64-bit]).

### Mobility

Animals were either synchronized by timed egg-lay (two replicates) or by hypochlorite treatment (one replicate) on RNAi control plates. When reaching adulthood, nematodes were transferred to RNAi plates. Every few days at regular intervals, plates were rocked in order to induce movement of animals and videos were subsequently recorded for one minute. Worms were transferred to fresh plates whenever necessary. At day 16 the vast majority of worms completely ceased movement, thus we did not include any later timepoints. Notably, we did not notice any obvious aversion behavior or elevated speed at young age upon *nsun-1* or *nsun-5* RNAi, which was previously shown to be present upon depletion of other components of the translational machinery (Melo and Ruvkun, 2012). Worm Lab version 4.1.1 was used to track individual animals and calculate the average speed.

### Lifespan assays

Lifespan measurement was conducted as previously described (Schosserer et al., 2015). For lifespan assays, 90 adults per condition were transferred to plates seeded with the respective RNAi bacteria (control, *nsun-1*, *nsun-5*). Wildtype worms were pre-synchronised on NGM plates seeded with UV-killed OP50 bacteria. 50 adult worms were transferred to NGM plates and allowed to lay eggs for 15 h; then the adult worms were removed. Synchronisation by timed egg lay was performed 72 h after the pre-synchronisation by transferring 350 gravid worms from the pre-synchronisation to fresh NGM plates seeded with RNAi control bacteria and allowed to lay eggs for four hours. After 68 h, 90 young adult worms per condition were placed on fresh NGM plates containing 5 mL NGM, 100 µL bacterial suspension and 50 μg FUDR. This day represents day 0 in the lifespan measurement. Worms were scored as “censored” or “dead” every two to four days. Nematodes were scored as “censored” if they had crawled off the plate, were missing or died due to other causes than ageing, such as gonad extrusion. Animals were transferred to fresh plates every 3–7 days depending on the availability of the bacterial food source. Lifespans were performed at 20°C. Kaplan-Meier survival curves were plotted and log-rank statistics were calculated.

### Thermotolerance

Thermotolerance was assessed as previously described (Vieira et al., 2017). Animals were synchronized by hypochlorite treatment and released eggs were transferred to NGM plates seeded with RNAi control bacteria and kept at 20°C. After 48 h, L4 animals were picked on RNAi control, *nsun-1* or *nsun-5* RNAi plates and exposed to RNAi for approximately three days (68 hours). Subsequently, plates were transferred to 35°C and scored every 1-2 h for survival.

### Body size

Worms were synchronized by hypochlorite treatment and incubated in liquid S-Basal medium overnight. On the following day, eggs/L1 were transferred to RNAi plates (RNAi control, *nsun-1* and *nsun-5* RNAi). Three days later, worms were transferred to agar pads and paralyzed using sodium azide and visualized using DIC microscopy (see above).

### Brood size analysis

Worms were synchronised by treatment with hypochlorite solution and incubated in S-Basal at room temperature overnight. L1 larvae were subsequently transferred to NGM plates seeded with RNAi control bacteria. After 48 h, L4 animals were transferred to individual wells of a 24-well plate seeded with the respective RNAi bacteria (HT115, *nsun-1*, *nsun-5*). Each well contained 1.5 mL of NGM agar and 3 μL of bacterial suspension (1:2 dilution in S-Basal). Worms were transferred to a new well every day for four consecutive days and total progeny of individual animals was counted. Per condition and experiment, five worms were analysed.

### Global protein synthesis by puromycin incorporation

Puromycin incorporation was measured as previously described (Tiku et al., 2018) with minor modifications. Heat-inactivated OP-50 (75°C, 40 min) were provided as food source during pulse labelling. As negative controls, RNAi control treated worms were used either without addition of puromycin or by pulse-labelling at 4°C instead of 20°C. Around 100 animals per condition were harvested for western blot analysis. Lysis was done directly in SDS loading dye (60 µM Tris/HCl pH 6.8, 2% SDS, 10% glycerol, 0.0125% bromophenol blue and 1.25% β-mercaptoethanol). Worms in SDS loading dye were homogenized with a pellet pestle for 1 min. Then, the samples were heated to 95°C and loaded on 4-15% Mini-PROTEAN^®^ TGX gels (BioRad) in Laemmli-Buffer (25 mM Tris, 250 mM glycine and 0.1% SDS). Protein bands were transferred to PVDF-membranes (Bio Rad) at 25 V and 1.3 A for 3 min. After blocking with 3% milk in PBS, the membrane was incubated overnight at 4°C with a mixture of anti-Histone H3 (Abcam ab1791, 1:4000) and anti-puromycin (Millipore 12D10, 1:10000). After washing and secondary antibody incubation (IRDye680RD and IRDye800CW, 1:10000), the membrane was scanned on the Odyssey Infrared Imager (LI-COR). Quantification of band intensities was performed in Image J (version 2.0.0-rc- 65/1.51w; Java 1.8.0_162 [64-bit]).

### Polysome profling

Two-day-old adult worms were used to generate polysome profiles as previously described (Rogers et al., 2011). One hundred microliter worm-pellet were homogenized on ice in 300 µL of solubilisation buffer (300 mM NaCl, 50 mM Tris-HCl (pH 8.0), 10 mM MgCl_2_, 1 mM EGTA, 200 µg/ml heparin, 400 U/ml RNAsin, 1.0 mM phenylmethylsulfonyl fluoride, 0.2 mg/ml cycloheximide, 1% Triton X-100, 0.1% sodium deoxycholate) using a pellet pestle. 700 µl additional solubilisation buffer were added, vortexed briefly, and placed on ice for 10 min before centrifugation at 20.000 g for 15 min at 4°C. Approximately 0.9 mL of the supernatant was applied to the top of a linear 10-50% sucrose gradient in high salt resolving buffer (140 mM NaCl, 25 mM Tris-HCl (pH 8.0), 10 mM MgCl_2_) and centrifuged in a Beckman SW41Ti rotor (Beckman Coulter, Fullerton, CA, USA) at 180.000 g for 90 min at 4°C.

### RNA Seq

Gradients were fractionated while continuously monitoring the absorbance at 260 nm. Trizol LS (Life Technologies) was immediately added to collected fractions and RNA was isolated following the manufacturer’s protocol. PolyA-selection, generation of a strand-specific cDNA library and sequencing on the HiSeq 4000 platform (Illumina) using the 50 bp SR mode was performed by GATC Biotech (Konstanz, Germany). At least 30 million reads were generated per sample.

FASTQ Trimmer by column (Galaxy Version 1.0.0) was used to remove the first 12 bases from the 5’ end of each read due to an obvious base bias in this region, as detected by FastQC (Galaxy Version 0.69). Filter by quality (Galaxy Version 1.0.0) was performed using a cut-off value of 20 and only reads with a maximum number of 8 bases with quality lower than the cut-off value were retained. RNA STAR (Galaxy Version 2.6.0b-1) was used to align reads to the WBcel235 reference genome using the default options. Aligned reads with a minimum alignment quality of 10 were counted using htseq-count (Galaxy Version 0.9.1).

Differential expression was analyzed using the DEseq2 package in R. The contrast

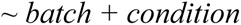

(batch = biological replicate, condition = sample description) was applied to compare the polysome fraction to the total RNA of either RNAi control or *nsun-1* RNAi treated samples. Afterwards, results were filtered in R to contain only protein-coding genes (according to ENSEMBL annotation), genes with detectable expression (base mean > 1), a fold change of > 2 (log2FC > 1) and an adjusted p-value of < 0.05. Vulcano plots were generated in R using the EnhancedVulcano package, labelling the top 5 up- and down-regulated genes respectively.

GO term enrichment using DAVID (version 6.7) was performed as previously described (Rollins et al., 2019). Only protein-coding genes with detectable expression (base mean > 1), a fold change of > 2 and an adjusted p-value of < 0.10 were considered. For visualization, only the broadest GO terms of the GOTERM_BP_FAT, GOTERM_MF_FAT and GOTERM_CC_FAT categories, which were still significantly enriched (FDR < 0.05), are shown while similar terms based on the same subset of genes but lower in hierarchy were manually removed. Full results are contained in the supplements. UTR characterization and RBP motif enrichment were performed as previously described (Rollins et al., 2019) using only protein-coding genes with detectable expression (base mean > 1), a fold change of > 2 and an adjusted p-value of < 0.10.

The raw and processed sequencing data are available from the Gene Expression Omnibus database (https://www.ncbi.nlm.nih.gov/geo) under accession GSE143618.

The R-script for analyzing RNA-seq data is provided as Supplemental Data File 5.

### RT-qPCR

Samples were collected by either transferring worms individually into 1.5 mL tubes or by washing them off NGM plates using S-Basal. After three washing steps with S-Basal, 300 μL TRIzol^®^ LS Reagent were added to approximately 100 µL residual S-Basal including worms. Subsequently, worms were homogenised with a pellet pestle for one minute, 600 μl TRIzol^®^ LS Reagent were added and the sample was vortexed for five minutes at room temperature. Total RNA was isolated using Direct-zol™ RNA MiniPrep Kit (Zymo) according to the instructions by the manufacturer. For cDNA synthesis, 500 ng RNA were converted into cDNA using the Applied Biosystems™ High-Capacity cDNA Reverse Transcription Kit (Thermo Fisher Scientific). cDNA was amplified from total RNA using random primers. RT-qPCR was performed on a Rotor-Gene Q (QIAGEN) using HOT FIREPol^®^ EvaGreen^®^ qPCR Mix. The absolute amounts of mRNAs were calculated by computing a standard curve and the resulting copy numbers were normalized to the housekeeping genes act-1 and tba-1. The following primers were used: <u>nsun-1</u>: 5’-TCGCCGAGATCCACAGAAAT-3’ (sense) and 5’-CCACGTTCATTCCACGGTTG-3’ (antisense); <u>nsun-2:</u> 5’-GCTTAAACGAGAGACGGGAGTT-3’ (sense) and 5’-CACCAGTATCCTGGGCGTG-3’ (antisense); <u>nsun-4:</u> 5’-TGTTGGATATGTGTGCGGCT-3’ (sense) and 5’-GCGTCCTTGCGTTTTAGGAC-3’ (antisense); <u>nsun-5:</u> 5’-GGCCAAGGAGAAAAGTGTG-3’ (sense) and 5’-GATCCACCGATATTCGCAT-3’ (antisense); <u>act-1:</u> 5’-CTACGAACTTCCTGACGGACAAG-3’ (sense) and 5’-CCGGCGGACTCCATACC-3’ (antisense) and <u>tba-1:</u> 5’-TCAACACTGCCATCGCCGCC-3’ (sense) and 5’-TCCAAGCGAGACCAGGCTTCAG-3’ (antisense).

For measuring mRNA expression during development, worms were synchronised by treatment with hypochlorite solution and the released eggs were subsequently transferred to four separate NGM plates seeded with UV-killed OP50 bacteria. Samples were taken from eggs immediately after bleaching, L1/L2 (20 h after bleaching), L3 (32 h after bleaching), L4 (46 h after bleaching) and young adults (60 h after bleaching).

### 3-D ribosome structure

The PyMOL Molecular Graphics System (Version 2.0) was used. The structure was modelled on the human 80S ribosome (PDB 6EK0).

### m^5^C detection by COBRA assay

NSUN-5 activity was measured by the COBRA assay as previously described (Adamla et al., 2019) using the following primers: 5’-GGGAGTAATTATGATTTTTCTAAGGTAG-3’ (sense) and 5’-ATAATAAATAAAAACAATAAAAATCTCACTAATCCATTCATACAC-3’ (antisense).

### HPLC analysis of m^5^C

13-15µg 26S purified on sucrose gradient were digested to nucleosides and analyzed by HPLC. Peaks elutes at 12 min and as a control, a commercial 5-methylcytidine (NM03720, CarboSynth) was used. For quantification of m^5^C peak area, the peak was normalized to either the peak eluting at 16 min (asterisk on the Figure), or to the peak eluting at 8 min (U), with similar results. The results are shown for normalization to the peak eluting at 16 min.

### Pre-rRNA processing analysis

For analysis of high–molecular weight RNA species, 3 µg total RNA was resolved on a denaturing agarose gel (6% formaldehyde/1.2% agarose) and migrated for 16 h at 65 volts. Agarose gels were transferred by capillarity onto Hybond-N+ membranes. The membrane was prehybridized for 1 h at 65°C in 50% formamide, 5x SSPE, 5x Denhardt’s solution, 1% SDS (w/v) and 200 µg/ml fish sperm DNA solution (Roche). The ^32^P-labeled oligonucleotide probe (LD2648 (ITS1): CACTCAACTGACCGTGAAGCCAGTCG; LD2649 (ITS2): GGACAAGATCAGTATGCCGAGACGCG) was added and incubated for 1 h at 65°C and then overnight at 37°C. For analysis of low molecular weight RNA species, northern blots were exposed to Fuji imaging plates (Fujifilm) and signals acquired with a Phosphorimager (FLA-7000; Fujifilm).

### Statistics and sample size estimation

No explicit power analysis was used. Sample sizes estimations were partially based on our own previous empirical experience with the respective assays, as well as the cited literature.

No systematic blinding of group allocation was used, but samples were always analysed in a random order. Nematodes were randomly assigned to the experimental groups. All lifespan, stress resistance and locomotion experiments were performed by at least two different operators.

Most experiments were performed in three independent experiments, unless stated otherwise in the figure legend. Independent experiments were always initiated at different days and thus always resemble different batches of nematodes. Some experiments (RNA isolation for RNA-seq, HPLC analysis of m^5^C, pre-rRNA processing analysis) were performed once with all frozen independent batches of nematodes to minimize technical variation. No outliers were detected or removed. Criteria for censoring animals for lifespan, stress resistance and locomotion experiments are indicated in the respective chapters.

Statistical tests used, exact values of N, definitions of center, methods of multiple test correction, and dispersion and precision measures are indicated in the respective figure legends. P-value thresholds were defined as *P < 0.05, **P < 0.01 and ***P < 0.001. For RNA-seq, statistical tests and p-value thresholds are explained in detail in the “RNA-seq” chapter.

## Acknowledgements

We are grateful to Tamás Barnabás Könye for technical assistance; the BOKU-VIBT Imaging Center for technical support with microscopy. This work was supported by the Austrian Science Fund (FWF) and Herzfelder’sche Familienstiftung [P30623 to M.S.], Hochschuljubiläumsstiftung der Stadt Wien [H-327123/2018 to M.S.], and the Austrian Science Fund (FWF) [I2514 to J.G.]. Research reported in this publication was supported by the James L. Boyer Fellowship at the MDI Biological Laboratory to M.S. Research conducted in the labs of A.N.R and J.A.R. was supported by an Institutional Development Award (IDeA) from the National Institute of General Medical Sciences of the National Institutes of Health under grant numbers P20GM103423 and P20GM104318. Research in the Lab of D.L.J.L. is supported by the Belgian Fonds de la Recherche Scientifique (F.R.S./FNRS), the Université Libre de Bruxelles (ULB), the Région Wallonne (DGO6) [grant RIBO*cancer* n°1810070], the Fonds Jean Brachet, the International Brachet Stiftung, and the Epitran COST action (CA16120). F.N. is a fellow of the international PhD programme “BioToP-Biomolecular Technology of Proteins”, funded by the Austrian Science Fund (FWF) [W1224 to J.G.]. Some strains were provided by the CGC, which is funded by NIH Office of Research Infrastructure Programs (P40 OD010440).

## Author Contributions

Planned experiments: C.H., T.L.K., J.A.R., J.G., A.N.R., D.L.J.L., M.S., performed experiments: C.H., T.L.K., I.S., A.S., S.S., M.S., analysed data: C.H., T.L.K., J.A.R., L.W., D.L.J.L., M.S., wrote manuscript: C.H., D.L.J.L., M.S., supervised the project: M.S.

## Figure Supplements

**Figure 1-figure supplement 1:**
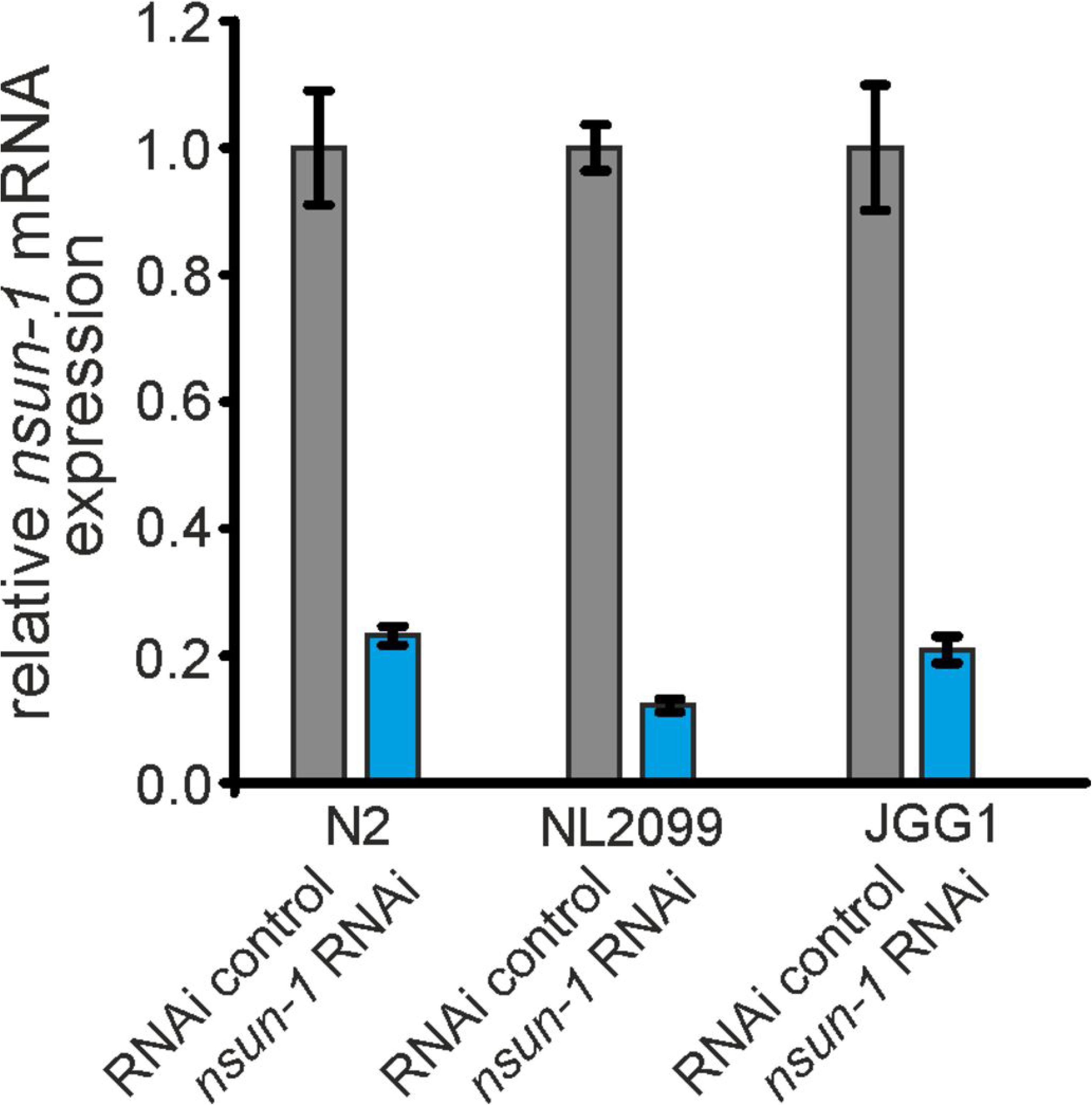
RNAi effectively depletes *nsun-1* in *C. elegans*. Quantification of *nsun-1* mRNA levels using RT-qPCR in N2, NL2099 and JGG1 nematode strains. Worms were subjected to control and *nsun-1* RNAi. *nsun-1* mRNA levels were decreased to approximately 20% in N2 and JGG1, as well as to 10% in the RNAi hypersensitive strain NL2099. *act-1* was used for normalization. Error bars represent standard deviation of four technical replicates. This experiment was repeated independently with similar outcome.

**Figure 3-figure supplement 1:**
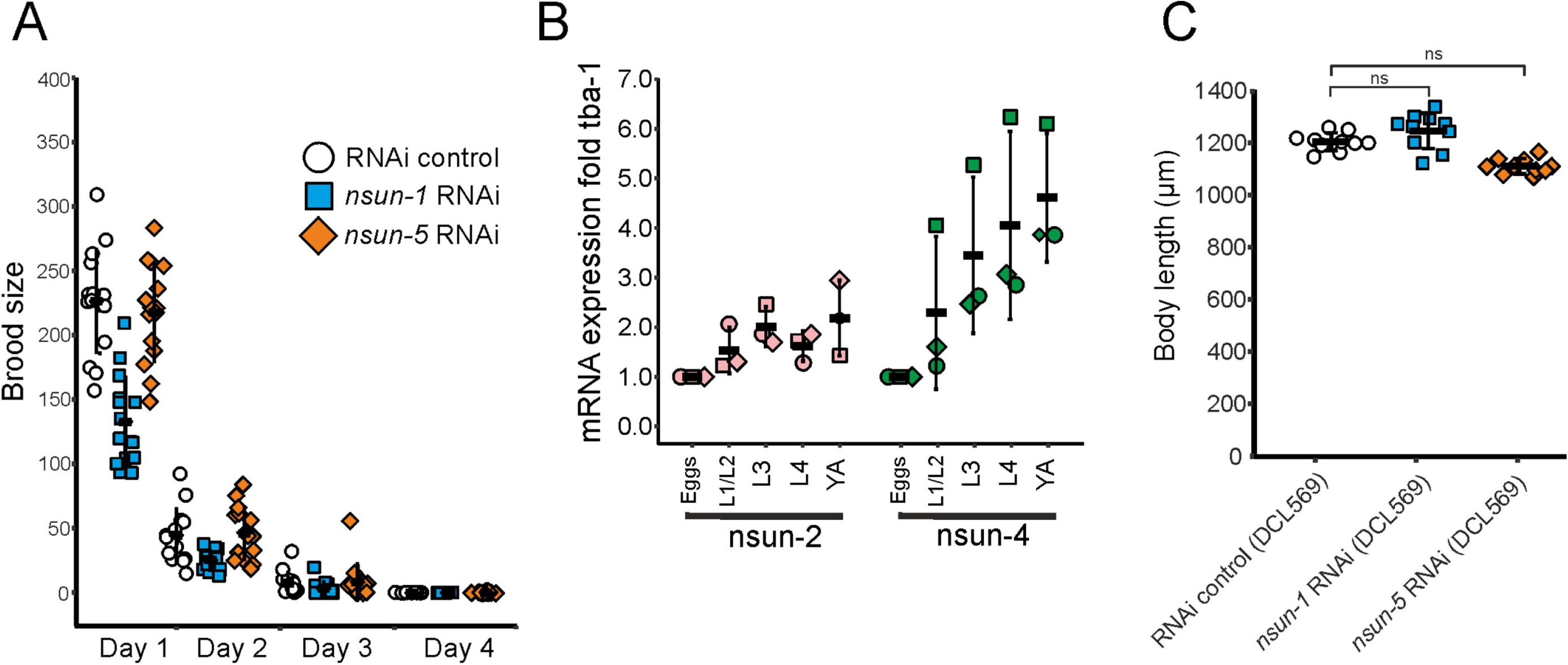
*nsun-1* depleted worms display impaired fecundity. **A,** Brood size analysis of adult-onset RNAi exposed animals. Eggs of individual worms were counted every day until day 4 of adulthood. Knockdown of *nsun-1*, but not *nsun-5*, inflicted a reduced brood size compared to control RNAi. Three pooled independent experiments are shown. n = 5 per condition and per experiment. Error bars indicate standard deviation. **B,** RT-qPCR analysis of developing wild-type animals (eggs, L1/L2 larvae, L3 larvae, L4 larvae and young adults) revealed enhanced mRNA expression of *nsun-2* and *nsun-4* during development. Three independent biological experiments are shown. *tba-1* was used for normalization. Error bars represent standard deviation. **C,** Body size of 1-2 day old adult worms of a germline-specific RNAi strain (DCL569) was measured. The body size of *nsun-1* and *nsun-5* RNAi treated animals was not changed compared to RNAi control. n(all conditions) = 10, one-way ANOVA with Dunnett’s post, α=0.05, not significant. Error bars represent standard deviation.

**Figure 4-figure supplement 1:**
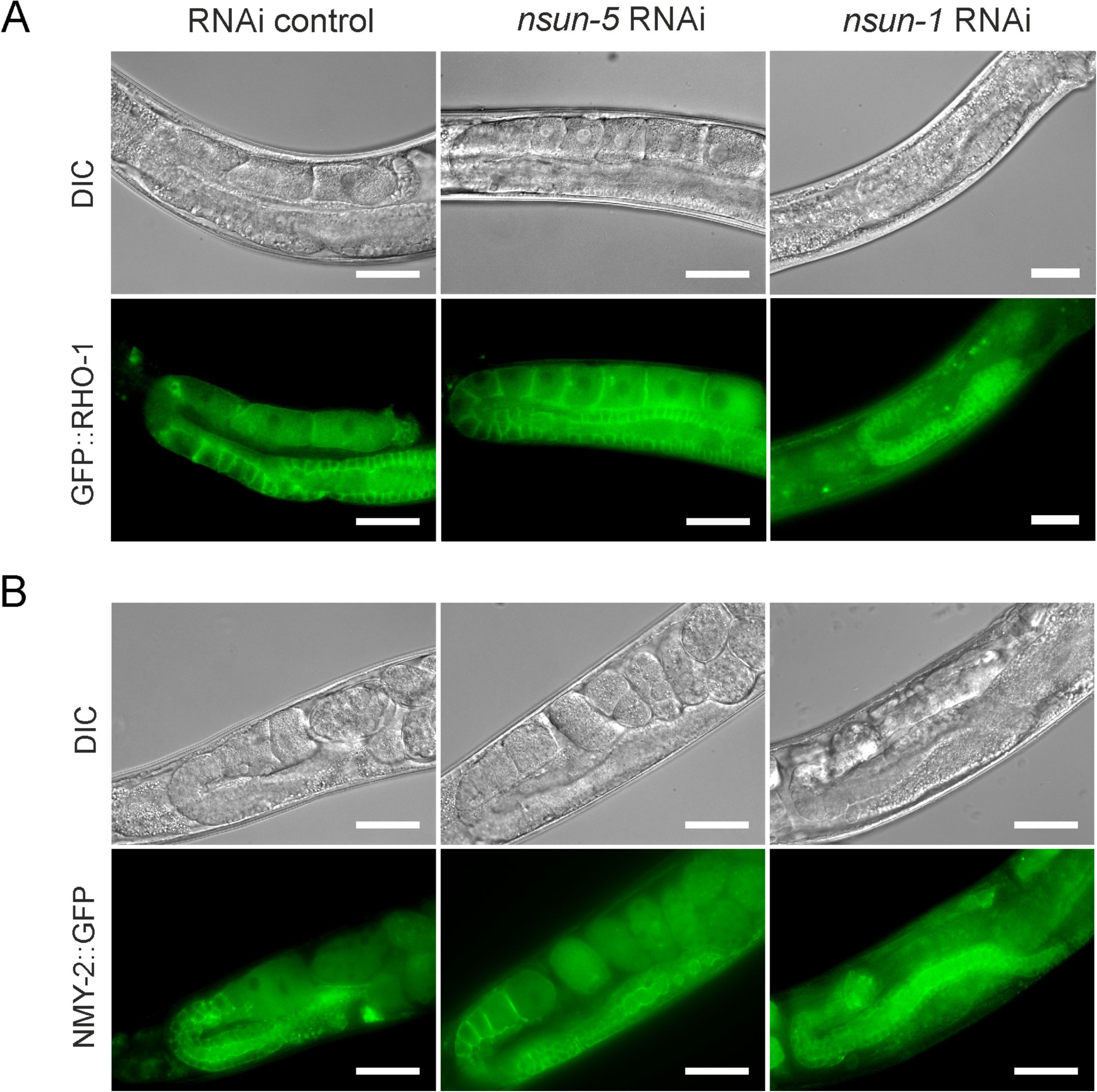
*nsun-1* but not *nsun-5* depletion inflicts a defect in oogenesis. **A, B,** Gonad specific expression of GFP::RHO-1 (SA115 strain) **(A)** and NMY-2::GFP (JJ1473 strain) **(B)** were used to visualize the morphology of the germline after *nsun-1* and *nsun-5* knockdown. Scale bar represents 40 µm. Oozyte maturation starting from the loop region was impaired in *nsun-1* RNAi, but not in *nsun-5* RNAi treated animals.

**Figure 4-figure supplement 2:**
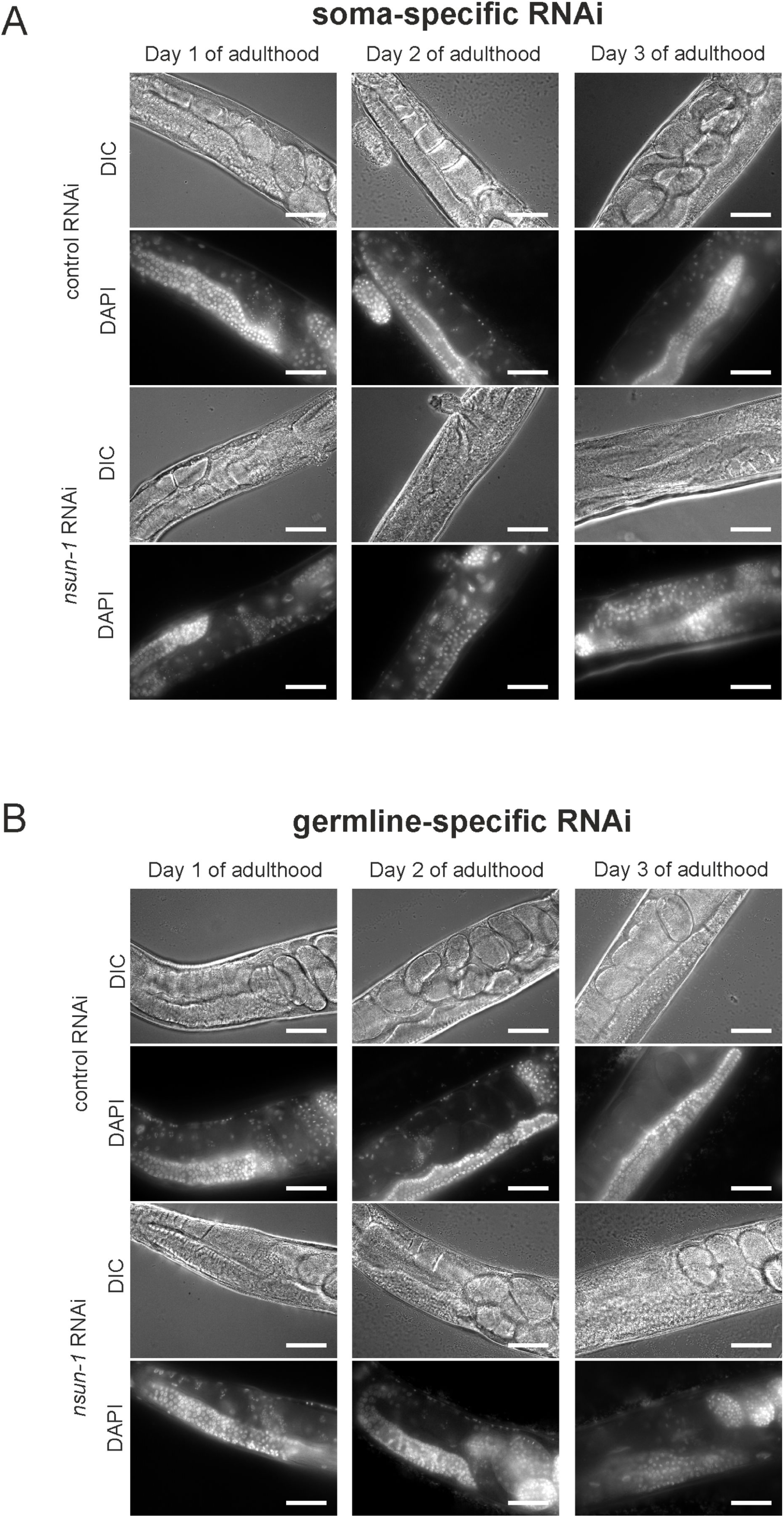
Somatic- but not germline specific nsun-1 depletion causes defective oogenesis. **A-B,** Soma- but not germline-specific *nsun-1* RNAi phenocopies altered gonad morphology of whole body *nsun-1* depletion. NL2550 was used for soma- **(A)** and NL2098 for germline-specific knockdown **(B)**. The gonad of 1-, 2- and 3-day old animals was imaged in DIC mode and nuclei were stained with DAPI following fixation. In the soma- specific strain maturating oocytes and embryos were observed only in RNAi control but not in *nsun-1* RNAi subjected worms. The germline-specific strain remained entirely unaffected. Scale bar represents 40 µm.

**Figure 5-figure supplement 1:**
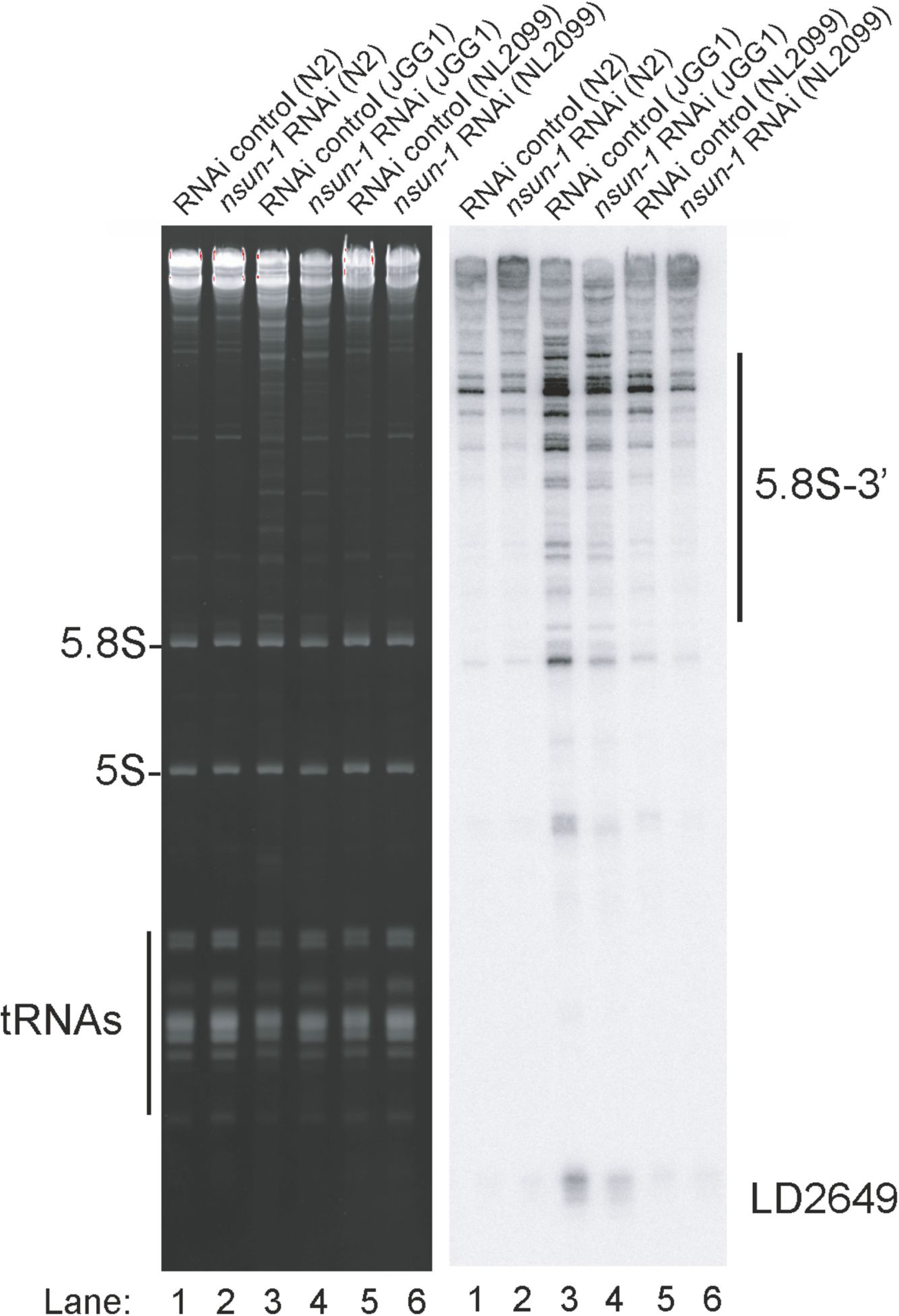
*nsun-5* but not *nsun-1* RNAi alters 5.8S rRNA maturation. In the absence of *nsun-5* (JGG1 strain, lane 3 and 4) the 3′-extended forms of 5.8S and short RNA degradation products accumulated. Upon co-depletion of *nsun-1* (lane 4), such an accumulation is partially suppressed. When comparing the 5.8S and 5S, as well as tRNAs, no change can be observed between control and *nsun-1* RNAi treated animals in the three different worm strains (N2, JGG1 and NL2099).

**Figure 6-figure supplement 1:**
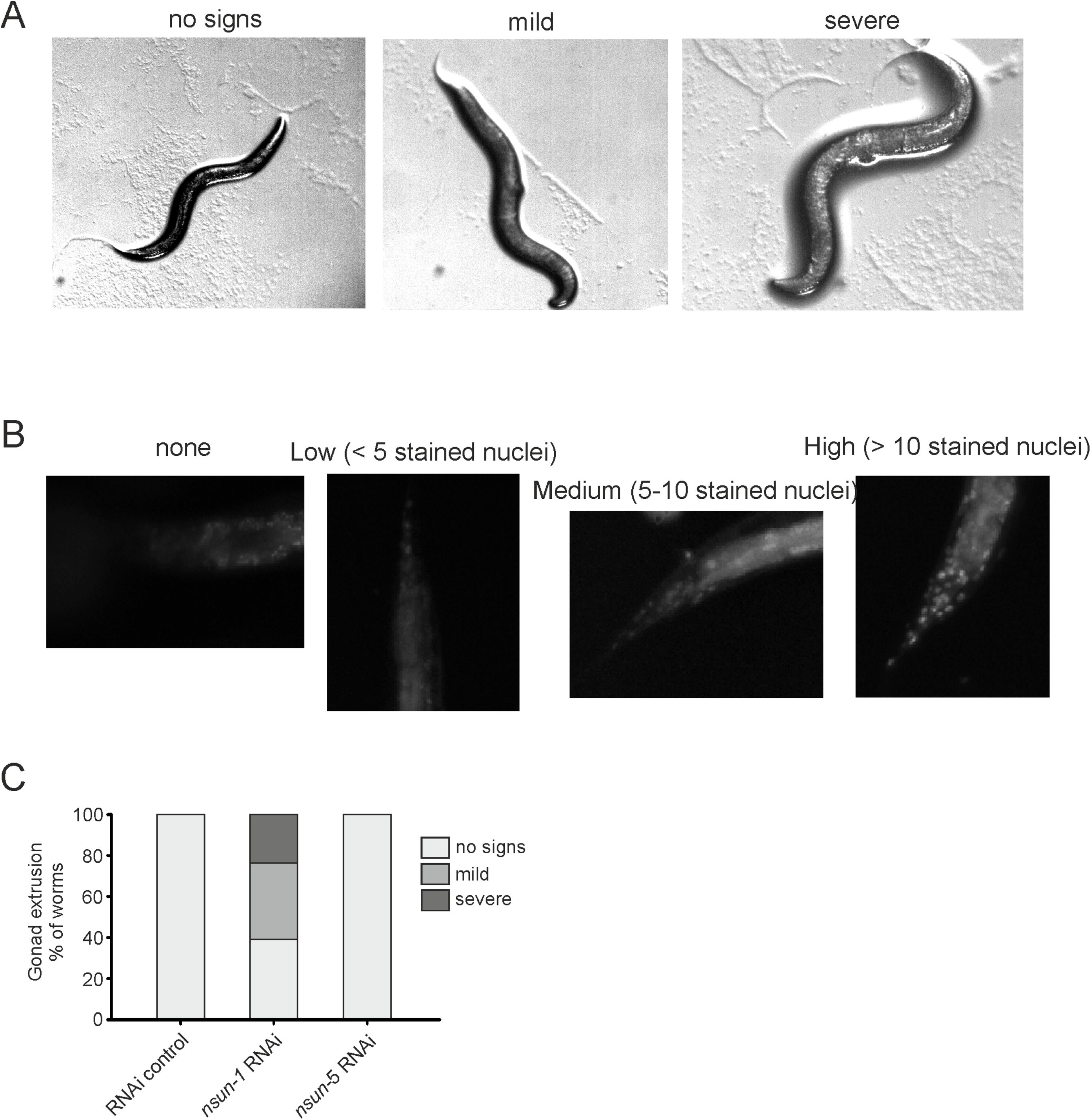
*nsun-1* depletion affects gonad integrity and barrier function. **A,** Knockdown of *nsun-1* increased the rate and severity of gonad extrusion. Mid-aged worms at day 8-9 of adulthood were classified into three categories according to the severance of gonad extrusion (‘no signs’, ‘mild’, ‘severe’, see arrowhead). Representative images of the categories are shown here. **B,** Reduced levels of *nsun-1* affected barrier function. Young adult animals were incubated in 1 µg/mL Hoechst 33342, which is membrane-permeable but cuticle-impermeable. Permeability was assessed by counting nuclear Hoechst staining in the tail region (see arrowhead). Young adult animals were classified into four categories (‘no staining’, ‘low (< 5 stained nuclei)’, ‘medium (5-10 stained nuclei)’, ‘high (>10 stained nuclei)’). Representative images of the different categories are shown here. **C,** Analysis of gonad extrusion including the results of *nsun-5* RNAi. The experiment was independently performed two times with similar outcome. One representative replicate is shown. n ≥ 50 animals per replicate.

## Source Data Files for Figures

Figure 2–Source Data 1: Summary of individual lifespan and thermotolerance experiments

## References

Adamla F, Rollins J, Newsom M, Snow S, Schosserer M, Heissenberger C, Horrocks J, Rogers AN, Ignatova Z. 2019. A Novel Caenorhabditis Elegans Proteinopathy Model Shows Changes in mRNA Translational Frameshifting During Aging. Cell Physiol Biochem 52:970–983. doi:10.33594/000000067

Ban N, Beckmann R, Cate JHD, Dinman JD, Dragon F, Ellis SR, Lafontaine DLJ, Lindahl L, Liljas A, Lipton JM, McAlear MA, Moore PB, Noller HF, Ortega J, Panse VG, Ramakrishnan V, Spahn CMT, Steitz TA, Tchorzewski M, Tollervey D, Warren AJ, Williamson JR, Wilson D, Yonath A, Yusupov M. 2014. A new system for naming ribosomal proteins. Curr Opin Struct Biol 24:165–9. doi:10.1016/j.sbi.2014.01.002

Bansal A, Zhu LJ, Yen K, Tissenbaum HA. 2015. Uncoupling lifespan and healthspan in *Caenorhabditis elegans* longevity mutants. Proc Natl Acad Sci 112:E277–E286. doi:10.1073/pnas.1412192112

Bantis A, Giannopoulos A, Gonidi M, Liossi A, Aggelonidou E, Petrakakou E, Athanassiades P, Athanassiadou P. 2004. Expression of p120, Ki-67 and PCNA as proliferation biomarkers in imprint smears of prostate carcinoma and their prognostic value. Cytopathology 15:25–31. doi:10.1046/j.0956-5507.2003.00090.x

Bar DZ, Charar C, Dorfman J, Yadid T, Tafforeau L, Lafontaine DLJ, Gruenbaum Y. 2016. Cell size and fat content of dietary-restricted Caenorhabditis elegans are regulated by ATX-2, an mTOR repressor. Proc Natl Acad Sci U S A 113:E4620–9. doi:10.1073/pnas.1512156113

Cheng JX, Chen L, Li Y, Cloe A, Yue M, Wei J, Watanabe KA, Shammo JM, Anastasi J, Shen QJ, Larson RA, He C, Le Beau MM, Vardiman JW. 2018. RNA cytosine methylation and methyltransferases mediate chromatin organization and 5-azacytidine response and resistance in leukaemia. Nat Commun 9:1163. doi:10.1038/s41467-018-03513-4

Chiocchetti A, Zhou J, Zhu H, Karl T, Haubenreisser O, Rinnerthaler M, Heeren G, Oender K, Bauer J, Hintner H, Breitenbach M, Breitenbach-Koller L. 2007. Ribosomal proteins Rpl10 and Rps6 are potent regulators of yeast replicative life span. Exp Gerontol 42:275–286. doi:10.1016/j.exger.2006.11.002

Cui W, Pizzollo J, Han Z, Marcho C, Zhang K, Mager J. 2016. Nop2 is required for mammalian preimplantation development. Mol Reprod Dev 83:124–31. doi:10.1002/mrd.22600

Curran SP, Ruvkun G. 2007. Lifespan regulation by evolutionarily conserved genes essential for viability. PLoS Genet 3:0479–0487. doi:10.1371/journal.pgen.0030056

Dong Y, Poulin G, Kanapink A, Bot N Le, Welchman DP, Zipperlen P, Ahringer J. 2003. Systematic functional analysis of the Caenorhabditis elegans genome using RNAi. Nature 421. doi:10.1038/nature01278

Ewald CY, Landis JN, Porter Abate J, Murphy CT, Blackwell TK. 2015. Dauer-independent insulin/IGF-1-signalling implicates collagen remodelling in longevity. Nature 519:97–101. doi:10.1038/nature14021

Genuth NR, Barna M. 2018. The Discovery of Ribosome Heterogeneity and Its Implications for Gene Regulation and Organismal Life. Mol Cell 71:364–374. doi:10.1016/j.molcel.2018.07.018

Gigova A, Duggimpudi S, Pollex T, Schaefer M, Koš M. 2014. A cluster of methylations in the domain IV of 25S rRNA is required for ribosome stability. RNA 20:1632–44. doi:10.1261/rna.043398.113

Hansen M, Taubert S, Crawford D, Libina N, Lee S-JJ, Kenyon C. 2007. Lifespan extension by conditions that inhibit translation in Caenorhabditis elegans. Aging Cell 6:95–110. doi:10.1111/j.1474-9726.2006.00267.x

Heissenberger C, Liendl L, Nagelreiter F, Gonskikh Y, Yang G, Stelzer EM, Krammer TL, Micutkova L, Vogt S, Kreil DP, Sekot G, Siena E, Poser I, Harreither E, Linder A, Ehret V, Helbich TH, Grillari-Voglauer R, Jansen-Dürr P, Koš M, Polacek N, Grillari J, Schosserer M. 2019. Loss of the ribosomal RNA methyltransferase NSUN5 impairs global protein synthesis and normal growth. Nucleic Acids Res 1–19. doi:10.1093/nar/gkz1043

Herovici C. 1963. [Picropolychrome: histological staining technic intended for the study of normal and pathological connective tissue]. Rev Fr Etud Clin Biol 8:88–9.

Hodgkin J, Barnes TM. 1991. More is not better: brood size and population growth in a self-fertilizing nematode. Proceedings Biol Sci 246:19–24. doi:10.1098/rspb.1991.0119

Hokii Y, Sasano Y, Sato M, Sakamoto H, Sakata K, Shingai R, Taneda A, Oka S, Himeno H, Muto A, Fujiwara T, Ushida C. 2010. A small nucleolar RNA functions in rRNA processing in Caenorhabditis elegans. Nucleic Acids Res 38:5909–18. doi:10.1093/nar/gkq335

Hsin H, Kenyon C. 1999. Signals from the reproductive system regulate the lifespan of C. elegans. Nature 399:362–6. doi:10.1038/20694

Janin M, Ortiz-Barahona V, de Moura MC, Martínez-Cardús A, Llinàs-Arias P, Soler M, Nachmani D, Pelletier J, Schumann U, Calleja-Cervantes ME, Moran S, Guil S, Bueno-Costa A, Piñeyro D, Perez-Salvia M, Rosselló-Tortella M, Piqué L, Bech-Serra JJ, De La Torre C, Vidal A, Martínez-Iniesta M, Martín-Tejera JF, Villanueva A, Arias A, Cuartas I, Aransay AM, La Madrid AM, Carcaboso AM, Santa-Maria V, Mora J, Fernandez AF, Fraga MF, Aldecoa I, Pedrosa L, Graus F, Vidal N, Martínez-Soler F, Tortosa A, Carrato C, Balañá C, Boudreau MW, Hergenrother PJ, Kötter P, Entian K-D, Hench J, Frank S, Mansouri S, Zadeh G, Dans PD, Orozco M, Thomas G, Blanco S, Seoane J, Preiss T, Pandolfi PP, Esteller M. 2019. Epigenetic loss of RNA-methyltransferase NSUN5 in glioma targets ribosomes to drive a stress adaptive translational program. Acta Neuropathol. doi:10.1007/s00401-019-02062-4

Kaeberlein M, Powers RW, Steffen KK, Westman EA, Hu D, Dang N, Kerr EO, Kirkland KT, Fields S, Kennedy BK. 2005. Regulation of Yeast Replicative Life Span by TOR and Sch9 in Response to Nutrients. Science (80- ) 310:1193–1196. doi:10.1126/science.1115535

Kapahi P. 2010. Protein synthesis and the antagonistic pleiotropy hypothesis of aging. Adv Exp Med Biol 694:30–7.

Kennedy S, Wang D, Ruvkun G. 2004. A conserved siRNA-degrading RNase negatively regulates RNA interference in C. elegans. Nature 427:645–9. doi:10.1038/nature02302

Kenyon C, Chang J, Gensch E, Rudner a, Tabtiang R, Jed AF, Kirk M, Davis, Kenyon C, Chang J, Gensch E, Rudner a, Tabtiang R. 1993. A C. elegans mutant that lives twice as long as wild type. Nature 366:461–464. doi:10.1038/366461a0

Kirkwood TB, Holliday R. 1979. The evolution of ageing and longevity. Proc R Soc Lond B Biol Sci 205:531–46.

Kumsta C, Hansen M. 2012. C. elegans rrf-1 mutations maintain RNAi efficiency in the soma in addition to the germline. PLoS One 7:e35428. doi:10.1371/journal.pone.0035428

Lee M-H, Schedl T. 2010. C. elegans star proteins, GLD-1 and ASD-2, regulate specific RNA targets to control development. Adv Exp Med Biol 693:106–22. doi:10.1007/978-1-4419-7005-3_8

Lithgow GJ, White TM, Hinerfeld DA, Johnson TE. 1994. Thermotolerance of a long-lived mutant of Caenorhabditis elegans. J Gerontol 49:B270–6. doi:10.1093/geronj/49.6.b270

Longman D, Johnstone IL, Cáceres JF. 2000. Functional characterization of SR and SR-related genes in Caenorhabditis elegans. EMBO J 19:1625–37. doi:10.1093/emboj/19.7.1625

Masoro EJ. 2005. Overview of caloric restriction and ageing. Mech Ageing Dev 126:913– 922. doi:10.1016/j.mad.2005.03.012

Melo J, Ruvkun G. 2012. Inactivation of conserved C. elegans genes engages pathogen- and xenobiotic-associated defenses. Cell 149:452–66. doi:10.1016/j.cell.2012.02.050

Natchiar SK, Myasnikov AG, Kratzat H, Hazemann I, Klaholz BP. 2017. Visualization of chemical modifications in the human 80S ribosome structure. Nature 551:472–477. doi:10.1038/nature24482

Ou H-L, Kim CS, Uszkoreit S, Wickström SA, Schumacher B. 2019. Somatic Niche Cells Regulate the CEP-1/p53-Mediated DNA Damage Response in Primordial Germ Cells. Dev Cell 50:167–183.e8. doi:10.1016/j.devcel.2019.06.012

Pan KZ, Palter JE, Rogers AN, Olsen A, Chen D, Lithgow GJ, Kapahi P. 2007. Inhibition of mRNA translation extends lifespan in Caenorhabditis elegans. Aging Cell 6:111–9. doi:10.1111/j.1474-9726.2006.00266.x

Pazdernik N, Schedl T. 2013. Introduction to germ cell development in Caenorhabditis elegans. Adv Exp Med Biol 757:1–16. doi:10.1007/978-1-4614-4015-4_1

Penzo M, Galbiati A, Treré D, Montanaro L. 2016. The importance of being (slightly) modified: The role of rRNA editing on gene expression control and its connections with cancer. Biochim Biophys Acta - Rev Cancer 1866:330–338. doi:10.1016/j.bbcan.2016.10.007

Rogers AN, Chen D, McColl G, Czerwieniec G, Felkey K, Gibson BW, Hubbard A, Melov S, Lithgow GJ, Kapahi P. 2011. Life span extension via eIF4G inhibition is mediated by posttranscriptional remodeling of stress response gene expression in C. elegans. Cell Metab 14:55–66. doi:10.1016/j.cmet.2011.05.010

Rollins JA, Howard AC, Dobbins SK, Washburn EH, Rogers AN. 2017. Assessing Health Span in Caenorhabditis elegans: Lessons From Short-Lived Mutants. J Gerontol A Biol Sci Med Sci 72:473–480. doi:10.1093/gerona/glw248

Rollins JA, Shaffer D, Snow SS, Kapahi P, Rogers AN. 2019. Dietary restriction induces posttranscriptional regulation of longevity genes. Life Sci alliance 2. doi:10.26508/lsa.201800281

Rual J-F, Ceron J, Koreth J, Hao T, Nicot A-S, Hirozane-Kishikawa T, Vandenhaute J, Orkin SH, Hill DE, van den Heuvel S, Vidal M. 2004. Toward improving Caenorhabditis elegans phenome mapping with an ORFeome-based RNAi library. Genome Res 14:2162–8. doi:10.1101/gr.2505604

Saijo Y, Sato G, Usui K, Sato M, Sagawa M, Kondo T, Minami Y, Nukiwa T. 2001. Expression of nucleolar protein p120 predicts poor prognosis in patients with stage I lung adenocarcinoma. Ann Oncol Off J Eur Soc Med Oncol 12:1121–5. doi:10.1023/a:1011617707999

Saijou E, Fujiwara T, Suzaki T, Inoue K, Sakamoto H. 2004. RBD-1, a nucleolar RNA-binding protein, is essential for Caenorhabditis elegans early development through 18S ribosomal RNA processing. Nucleic Acids Res 32:1028–1036. doi:10.1093/nar/gkh264

Schosserer M, Minois N, Angerer TB, Amring M, Dellago H, Harreither E, Calle-Perez A, Pircher A, Gerstl MP, Pfeifenberger S, Brandl C, Sonntagbauer M, Kriegner A, Linder A, Weinhäusel A, Mohr T, Steiger M, Mattanovich D, Rinnerthaler M, Karl T, Sharma S, Entian K-D, Kos M, Breitenbach M, Wilson IBH, Polacek N, Grillari-Voglauer R, Breitenbach-Koller L, Grillari J. 2015. Methylation of ribosomal RNA by NSUN5 is a conserved mechanism modulating organismal lifespan. Nat Commun 6:6158. doi:10.1038/ncomms7158

Sharma S, Lafontaine DLJJ. 2015. ‘View From A Bridge’: A New Perspective on Eukaryotic rRNA Base Modification. Trends Biochem Sci 40:560–575. doi:10.1016/j.tibs.2015.07.008

Sharma S, Langhendries J-L, Watzinger P, Kötter P, Entian K-D, Lafontaine DLJ. 2015. Yeast Kre33 and human NAT10 are conserved 18S rRNA cytosine acetyltransferases that modify tRNAs assisted by the adaptor Tan1/THUMPD1. Nucleic Acids Res 43:2242–58. doi:10.1093/nar/gkv075

Sharma S, Yang J, Watzinger P, Kötter P, Entian K-D. 2013. Yeast Nop2 and Rcm1 methylate C2870 and C2278 of the 25S rRNA, respectively. Nucleic Acids Res 1–15. doi:10.1093/nar/gkt679

Sijen T, Fleenor J, Simmer F, Thijssen KL, Parrish S, Timmons L, Plasterk RH, Fire A. 2001. On the role of RNA amplification in dsRNA-triggered gene silencing. Cell 107:465–76. doi:10.1016/s0092-8674(01)00576-1

Simsek D, Barna M. 2017. An emerging role for the ribosome as a nexus for post-translational modifications. Curr Opin Cell Biol 45:92–101. doi:10.1016/j.ceb.2017.02.010

Sleiman S, Dragon F. 2019. Recent Advances on the Structure and Function of RNA Acetyltransferase Kre33/NAT10. Cells 8:1035. doi:10.3390/cells8091035

Sloan KE, Warda AS, Sharma S, Entian K-D, Lafontaine DLJ, Bohnsack MT. 2017. Tuning the ribosome: The influence of rRNA modification on eukaryotic ribosome biogenesis and function. RNA Biol 14:1138–1152. doi:10.1080/15476286.2016.1259781

Syntichaki P, Troulinaki K, Tavernarakis N. 2007. eIF4E function in somatic cells modulates ageing in Caenorhabditis elegans. Nature 445:922–6. doi:10.1038/nature05603

Teuscher AC, Statzer C, Pantasis S, Bordoli MR, Ewald CY. 2019. Assessing Collagen Deposition During Aging in Mammalian Tissue and in Caenorhabditis elegans. Methods Mol Biol 1944:169–188. doi:10.1007/978-1-4939-9095-5_13

Tijsterman M, Okihara KL, Thijssen K, Plasterk RHA. 2002. PPW-1, a PAZ/PIWI protein required for efficient germline RNAi, is defective in a natural isolate of C. elegans. Curr Biol 12:1535–40. doi:10.1016/s0960-9822(02)01110-7

Tiku V, Kew C, Mehrotra P, Ganesan R, Robinson N, Antebi A. 2018. Nucleolar fibrillarin is an evolutionarily conserved regulator of bacterial pathogen resistance. Nat Commun 9:1–10. doi:10.1038/s41467-018-06051-1

Trixl L, Lusser A. 2019. The dynamic RNA modification 5-methylcytosine and its emerging role as an epitranscriptomic mark. Wiley Interdiscip Rev RNA 10:e1510. doi:10.1002/wrna.1510

Tushev G, Glock C, Heumüller M, Biever A, Jovanovic M, Schuman EM. 2018. Alternative 3’ UTRs Modify the Localization, Regulatory Potential, Stability, and Plasticity of mRNAs in Neuronal Compartments. Neuron 98:495–511.e6. doi:10.1016/j.neuron.2018.03.030

Vieira N, Bessa C, Rodrigues AJ, Marques P, Chan FY, de Carvalho AX, Correia-Neves M, Sousa N. 2017. Sorting nexin 3 mutation impairs development and neuronal function in Caenorhabditis elegans. Cell Mol Life Sci 1–18. doi:10.1007/s00018-017-2719-2

Voutev R, Killian DJ, Ahn JH, Hubbard EJA. 2006. Alterations in ribosome biogenesis cause specific defects in C. elegans hermaphrodite gonadogenesis. Dev Biol 298:45–58. doi:10.1016/j.ydbio.2006.06.011

Warnecke PM, Stirzaker C, Song J, Grunau C, Melki JR, Clark SJ. 2002. Identification and resolution of artifacts in bisulfite sequencing. Methods 27:101–7. doi:10.1016/s1046-2023(02)00060-9

Zou L, Wu D, Zang X, Wang Z, Wu Z, Chen D. 2019. Construction of a germline-specific RNAi tool in C. elegans. Sci Rep 9:2354. doi:10.1038/s41598-019-38950-8

